# Vagus nerve stimulation accelerates motor learning through cholinergic modulation

**DOI:** 10.1101/2021.10.28.466306

**Authors:** Spencer Bowles, Jordan Hickman, Xiaoyu Peng, W. Ryan Williamson, Rongchen Huang, Kayden Washington, Dane Donegan, Cristin G Welle

## Abstract

Vagus nerve stimulation (VNS) is a neuromodulation therapy for a broad and rapidly expanding set of neurologic conditions. Classically used to treat epilepsy and depression, VNS has recently received FDA approval for stroke rehabilitation and is under preclinical and clinical investigation for other neurologic indications. Despite benefits across a diverse range of neurological disorders, the mechanism through which VNS influences central nervous system circuitry is not well described, limiting therapeutic optimization. A deeper understanding of the influence of VNS on neural circuits and activity is needed to maximize the use of VNS therapy across a broad range of neurologic conditions.

To investigate how VNS can influence the neurons and circuits that underlie behavior, we paired VNS with upper limb movement in mice learning a skilled motor task. We leveraged genetic tools to perform optogenetic circuit dissection, as well as longitudinal *in vivo* imaging of calcium activity in cortical neurons to understand the effect of VNS on neural function. We found that VNS robustly enhanced motor learning when temporally paired with successful movement outcome, while randomly applied VNS impaired learning. This suggests that temporally-precise VNS may act through augmenting outcome cues, such as reinforcement signals. Within motor cortex, VNS paired with movement outcome selectively modulates the neural population that represents outcome, but not other movement-related neurons, across both acute and behaviorally-relevant timescales. Phasic cholinergic signaling from basal forebrain is required both for VNS-driven improvements in motor learning and the effects on neural activity in M1. These results indicate that VNS enhances motor learning through precisely-timed phasic cholinergic signaling to reinforce outcome, resulting in the recruitment of specific, behaviorally-relevant cortical circuits. A deeper understanding of the mechanisms of VNS on neurons, circuits and behavior provides new opportunities to optimize VNS to treat neurologic conditions.

## Introduction

Vagus Nerve Stimulation (VNS) is currently used in clinical care to treat epilepsy^1,2^ and depression^3^, but novel stimulation paradigms are being explored to treat a broad and growing range of neurologic injuries^4–6^. Recently, VNS temporally paired with motor rehabilitation was approved by the FDA for the treatment of motor deficits associated with stroke^6^. Preclinical and early clinical studies suggest that other paired-VNS paradigms can accelerate functional recovery from multiple neurologic conditions, including spinal cord injury, peripheral nerve injury and traumatic brain injury^7–9^. Despite the wide-ranging etiology of these conditions, the therapeutic model is similar: VNS is paired with a relevant rehabilitation protocol. It is hypothesized that this precise timing of stimulation drives targeted circuit plasticity for recovery from injury^5,10^. Yet, the lack of a clear circuit mechanism limits optimization of VNS therapy to treat neurologic injury.

The circuitry that mediates VNS effects on central nervous system plasticity remains poorly understood. Vagus nerve afferents terminate in the brainstem nucleus tractus solitarius (NTS)^11^, which in turn activates many subcortical and cortical brain regions, including major subcortical neuromodulatory nuclei^12–14^. Lesions of major neuromodulatory centers, including the cholinergic basal forebrain (BF), limit both VNS-driven cortical map plasticity^15,16^ and functional rehabilitation after peripheral nerve damage^9^. In addition, the cholinergic basal forebrain has been indicated as necessary for motor learning^17^, and phasic cholinergic signals are thought to play critical roles in reinforcement learning and outcome representation^18,19^. Together, these data suggest a possible role for phasic cholinergic signaling in mediating the effects of VNS-driven learning.

Paired VNS drives cortical map expansion specific to the associated sensory or motor representation. For instance, VNS paired with a forelimb movement expands the forelimb cortical representation^20^ while VNS paired with an auditory tone expands the cortical representation of that tone^21^. However, map expansion is delayed relative to changes in behavior, and does not always correlate to improved performance^22^. To achieve the improvements in motor and sensory learning, VNS must also influence specific neural activity and plasticity on shorter, behaviorally-relevant timescales. Electrophysiological^23–25^ and recent *in vivo* imaging studies have identified broad, excitatory effects of VNS across multiple cortical regions^14^. Yet, this non-specific alteration in excitatory drive cannot account for the selectivity of paired-VNS stimulation, which requires a specific refinement of relevant cortical circuits^8^.

To understand the mechanisms by which VNS can selectively modulate neural circuits to optimally enhance motor behavior, we compared the effect of VNS timing on skilled reach learning in mice and probed the underlying circuit using optogenetic cholinergic circuit manipulation, kinematic analysis, and *in vivo* calcium imaging in the motor cortex. Paired-enhanced skilled reach learning, but only when applied after a successful reach (Success VNS). Improved reach performance was explained by accelerated consolidation of reach trajectory onto an expert trajectory, indicating earlier and more effective motor learning. Cholinergic neural activity in the BF was required for the effects of VNS on motor learning and reach kinematics. VNS altered specific neural populations relevant to outcome representation in the primary motor cortex, and the effects of VNS in M1 were mitigated by cholinergic antagonists. These results indicate that VNS enhances motor learning through precisely-timed phasic cholinergic signaling to reinforce outcome, resulting in the recruitment of specific, behaviorally-relevant cortical circuits.

## Results

### VNS enhances skilled motor learning when paired with successful task outcome

To induce motor rehabilitation and cortical plasticity, VNS must be paired with movment^20,26^, yet an optimal pairing protocol has not been identified^8^. To determine the optimal timing of VNS during skilled motor learning, we applied multiple VNS pairing protocols as mice learned a skilled forelimb reaching task^27^. Using a newly developed chronic VNS approach for mice^28^, we implanted a microfabricated stimulation cuff on the left cervical vagus nerve (**Fig. 1a**), connected to a skull-mounted headcap. Mice were trained to perform a the skilled reach task, where they learn to reach through a slit to grab a food pellet off a post, for 14 days (**Fig. 1b,c**,).

**Figure 1.**
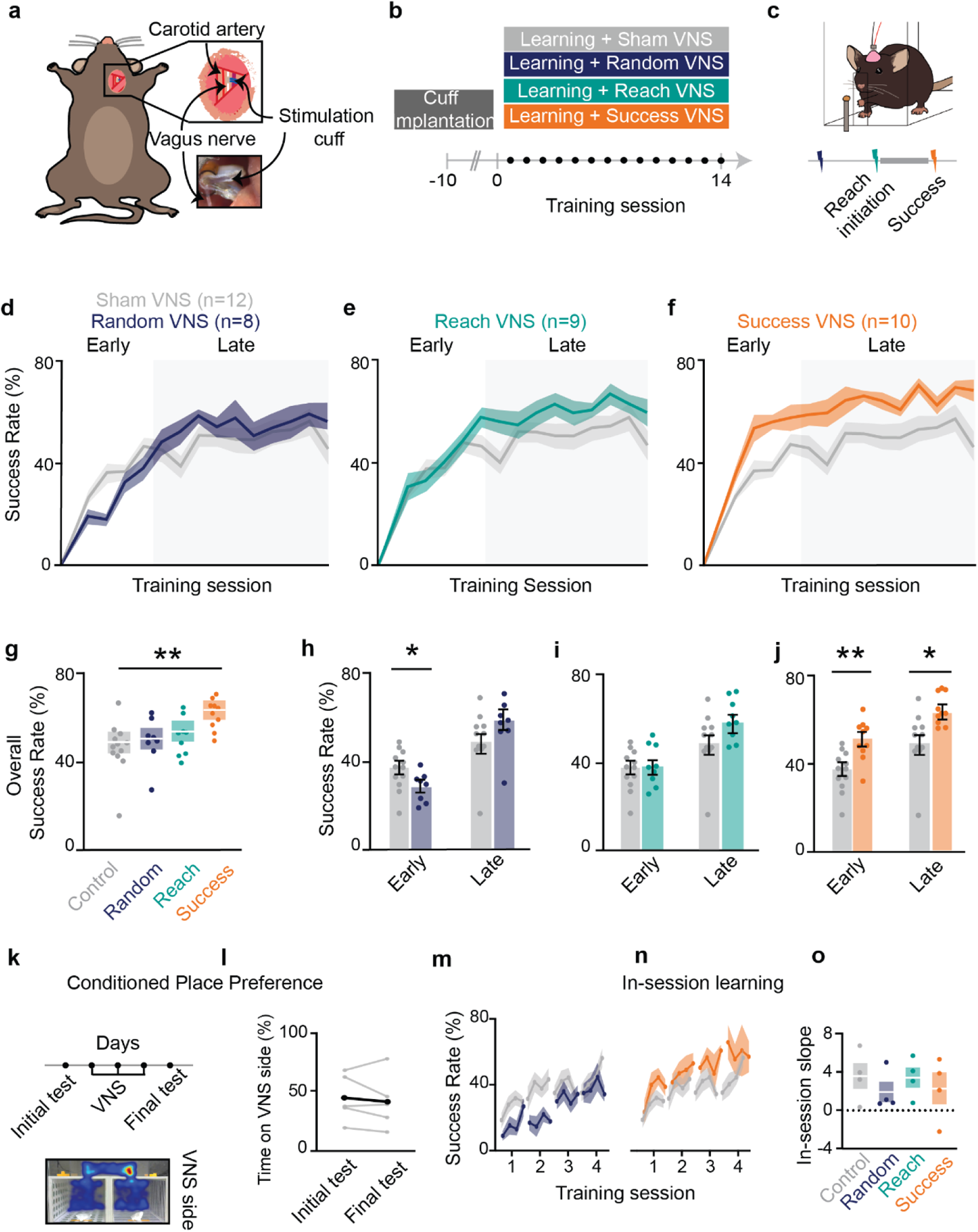
VNS modulates forelimb reach learning and requires temporally specific stimulation. **a,** VNS surgical approach **b,** Behavior timeline **c,** Stimulation protocol, with Reach and Success VNS applied before and after reach, respectively. **d-f,** Random VNS, Reach VNS, and Success VNS success rate across 14 sessions of training. **g,** Comparison of mean performance across all days between control and stimulated groups (Success VNS: p=0.0065, f=9.24, Random VNS: p>0.05, Reach VNS: p>0.05, REML). Shaded boxes denote s.e.m. **h,** Comparison of mean success rate for control and Random VNS mice during early (p=0.028, f=7.07, Student T test) and late learning (p>0.05). **i,** Comparison of mean success rate for control and Reach VNS mice during early and late learning (p>0.05). **j,** Comparison of mean success rate for control and Success VNS mice during early (p=0.0031) and late learning (p=0.0126). **k&l,** VNS mice performed a conditioned place preference test after 3 days being stimulated in one of two distinct rooms. **m&n,** In-session learning trajectories for each group. **o,** Comparison of within session learning between all groups across 4 days of learning. *p < 0.05, **p < 0.01, ***p< 0.001 bars and error bars represent the mean ± s.e.m.

We explored three possible mechanisms by which VNS could influence motor learning: arousal, spike-timing dependent plasticity and reinforcement (**Fig. 1b,c, see Methods for additional detail**). To test if VNS drives plasticity by increasing widespread cortical excitation and arousal^14,29,30^, VNS was applied at pseudo-random intervals (Random VNS) during the 20-minute training session. Alternatively, to determine if VNS acts through modulation of short-term attention or by influencing spike timing dependent plasticity^31^, VNS was applied at the initiation of a subset of reach movements (Reach VNS). To explore if VNS may augment reward or reinforcement related to movement outcome^32,33^, a third cohort received VNS after successful reach completion (Success VNS). The surgical control cohort was implanted with stimulation cuffs and connected to a stimulation isolation unit that was turned ‘off’ (Sham VNS). Current amplitude was consistent across groups and the number of stimulation trains delivered during training did not correlate with reach success rate (**Supp. Fig.1a-c**).

Animals in all cohorts learned to perform the skilled reach task (Sham VNS: p=0.0001, Random VNS: p=0.0001, Reach VNS: p=0.0002, Success VNS: p=0.001; **Supp. Fig. 2b-e**). Neither Random nor Reach VNS altered the success rate of the animals relative to Sham VNS (Random VNS: 47.4 ± 3.9%; Reach VNS: 53.6 ± 3.5%; Sham VNS: 46.3 ± 3.2%; **Fig. 1d,e,g**). However, Success VNS improved the overall success rate compared to Sham VNS (59.2 ± 3.1% vs 46.3 ± 3.2%; **Fig. 1f,g**), demonstrating for the first time that paired VNS can enhance motor learning in healthy animals.

**Figure 2.**
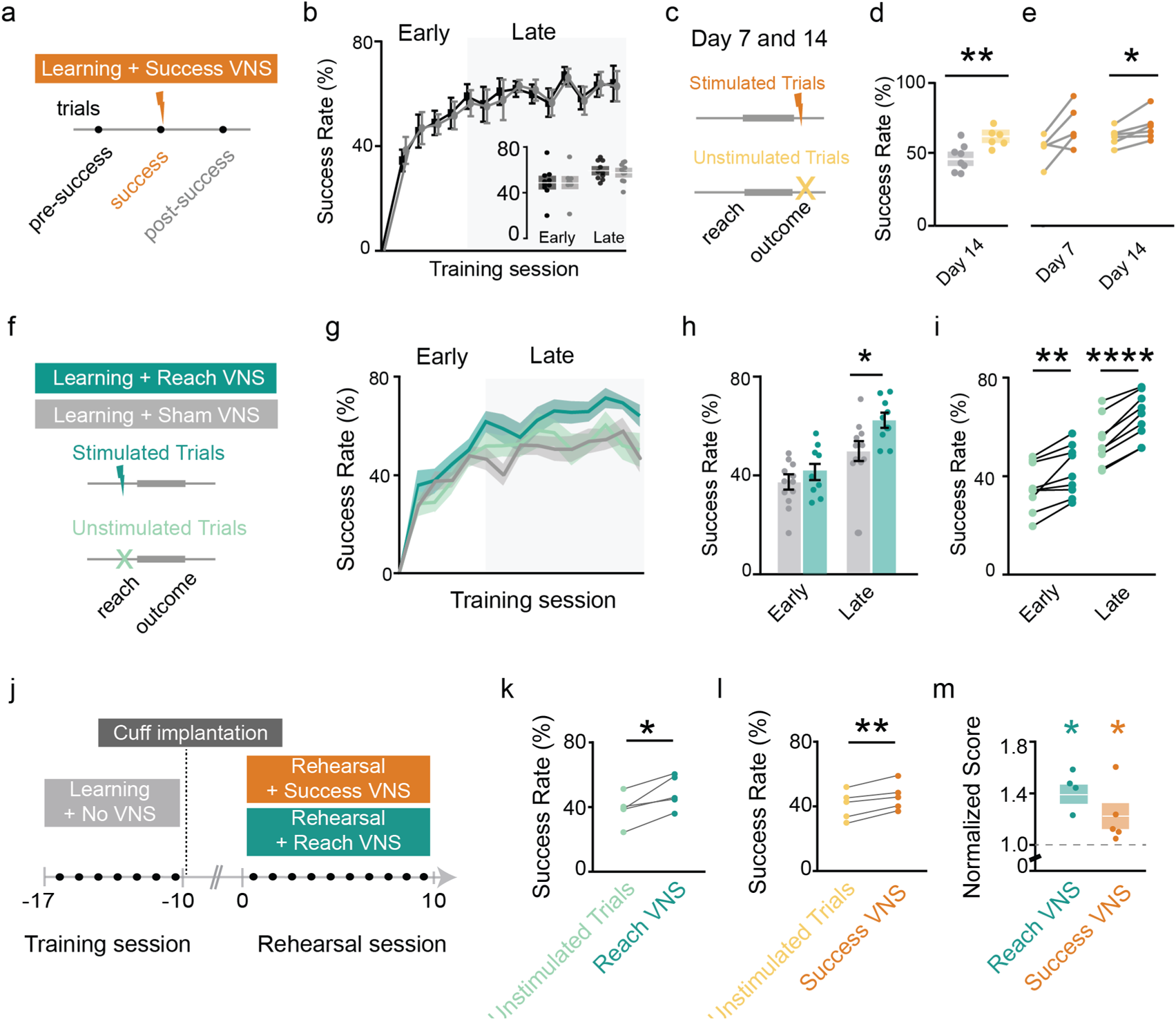
VNS improves success rate within sessions and in learned mouse during rehearsal of forelimb reach task. **a,** Trials before and after a stimulated success are investigated in success VNS. **b,** Comparison of success rate for reaches preceding or following stimulated reach. **c,** For Success VNS, day 7 and 14 trials are divided into equal blocks of unstimulated and stimulated trials. **d,** Comparison of unstimulated (light orange) and sham (grey) trials on day 14 (p=0.0059, t=3.34, Student’s T test). **e,** Comparison of stimulated (dark orange) and unstimulated (light orange) trials on days 7 & 14 (Day 7: p>0.05; Day 14: p=0.0499, t=2.57, Ratio paired t test). **f,** Reach VNS schematic. **g,** Success rates across learning training sessions for Sham (grey), Stimulated Reach VNS (dark green), and Unstimulated Reach VNS (light green). **h,** Comparison of Stimulated Reach VNS and Sham VNS trials in early (p>0.05) and late learning (p=0.024, t=2.47, Student’s T test). **i,** Comparison between stimulated Reach VNS and unstimulated Reach VNS trails in early (p=0.004, t=3.98, paired t test) and late learning (p<0.0001, t=10.08, paired t test). **j,** Success VNS and Reach VNS applied during rehearsal of reach task in trained mice. **k,** Stimulated Reach VNS trials improve success rate during rehearsal (p=0.015, t=4.058, paired t test). **l,** Stimulated Success VNS trials improve success rate during rehearsal (p=0.005, t=5.62, paired t test). **m,** Normalized improvement of Stimulated Reach VNS (p=0.047, t=3.56) and stimulated Success VNS trials (p=0.028, t=4.16) compared to unstimulated trials (Sidak’s multiple comparison’s test in RM one-way ANOVA). *p < 0.05, **p < 0.01, ***p < 0.001; bars and error bars represent the mean ± s.e.m.

Prior work on learning of a skilled reach suggests a multiphasic approach to learning, with distinct early and late learning phases^34,35^. Yet the timing of early to late transition has not been empirically demonstrated. Using a Weibull growth curve nonlinear model of the control learning curve, we identified an inflection point (55.49% ± 6.81) to determine early learning (days 1-4) and late learning (days 5-14; **Supp. Fig. 2a**). We next examined if VNS exerted distinct effects during different learning stages. Despite having no effect on the overall success rate, Random VNS impaired early learning (27.3 ± 7.3% vs. 37.4 ± 8.9%), but performance recovered during late learning (**Fig. 1h**). Reach VNS had no influence on success rates at any phase of learning (**Fig. 1i**). Success VNS increased success rates during the early and late phases (Early: 50.6 ± 9.4% vs. 37.4 ± 8.9%; Late: 63.6 ± 6.9 vs. 49.6 ± 13.2%; **Fig. 1j**). These data suggest that VNS paired with a successful outcome accelerates learning and increases the final proficiency of a forelimb task, while random application of stimulation temporarily impairs learning during early learning.

As only Success VNS enhanced motor learning, this indicated that VNS act through mechanisms paired with successful outcomes, likely reward or reinforcement. To determine if VNS serves as a rewarding or aversive stimulus^36^, we used a well-documented behavioral assessment of reward, the conditioned place preference test (CPP) (**Fig. 1k**). Implanted mice were introduced to two rooms with distinct visual and olfactory cues, with VNS applied in only one room for several days. On the final day probe session, mice spent equal time in the conditioned (stimulated) room as they did in their initial naïve session (44.8 ± 2.0% and 41.4 ± 2.3%; **Fig. 1l**), indicating that VNS is not inherently rewarding or aversive. Together, these results suggest that Success VNS may act by augmenting reinforcement cues, but not serve as a rewarding stimulus.

To determine if VNS alters within-session learning or between-session learning^37^, training session data was grouped into 4 blocks of 5 trials each (**Fig. 1m,n**), and the within-session learning slope was quantified over the first 4 days (**Fig. 1o**). The within-session learning slope was not significantly different between conditions, suggesting that Success VNS likely enhances between-session learning.

### VNS confers short-term performance benefits during the execution of learned tasks

Despite a lack of evidence for within-session learning, we wanted to confirm the behavioral results were due to learning, and not short-term modulation of attention. Since Success VNS is applied after reach outcome, the success rate of trials that immediately follow a stimulation (post-success) were compared to those immediately prior (pre-success). We found no effect of VNS on the success rate of trials following stimulation (**Fig. 2a,b**). We next compared the response to Success VNS to an unstimulated probe trial block included on days 7 and 14 (**Fig. 2c**). The success rate for unstimulated trials on Day 14 was greater than Sham VNS (61.9 ± 6.3% vs. 46.6 ± 9.2%; **Fig. 2d**), suggesting that Success VNS led to stimulation-independent, lasting learning. Yet, success rate during stimulated blocks was greater than unstimulated blocks on day 14 (69.7 ± 10.2% and 61.9 ± 6.3%), but not day 7 (**Fig. 2e**), implying a short-term performance benefit that emerges during late learning.

To further explore potential short-term benefits of VNS, we took advantage of the Reach VNS trial design, which allowed subgroup analysis of only stimulated or unstimulated reaches (**Fig. 2f**). The success rate of stimulated trials is greater than sham during late learning (64.0 ± 9.4% vs. 50.8 ± 13.7%; **Fig. 2h**), while unstimulated trials are not different from sham. Paired analysis for individual animals shows a higher success rate for stimulated trials compared to unstimulated trials in both the early and late phase (Early: 41.9 ± 10.2% vs. 35.3 ± 9.7%; Late: 64.0 ± 9.4% vs. 53.7 ± 9.5%; **Fig. 2i**). Taken together, Reach VNS provides a short-term performance boost for stimulated over unstimulated trials throughout learning. Similar to the results from Success VNS (**Fig. 2e**), Reach VNS most effectively modulates short-term performance for motor skills during late learning.

The prior results indicate the VNS can confer short-term performance benefit during late learning. To explore if this generalizes to tasks that are already known (learned without VNS), we applied paired VNS to animals already proficient in the skilled reaching task (**Fig. 2j**). Both Success VNS (**Fig. 2l,m**) and Reach VNS (**Fig. 2k,m**), delivered on alternate days for 10 days, improved performance over trial blocks without VNS (Success VNS: 46.0 ± 8.5% vs. 40.8 ± 8.9%; Reach VNS: 49.0 ± 10.2% vs. 38.8 ± 9.5%), confirming that either pairing protocol is sufficient to modulate the short-term performance of a known task. Together, this demonstrates that VNS confers short-term enhancement to performance of known motor skills.

### VNS drives neural activity in the basal forebrain (BF)

Cholinergic neuromodulation is associated with reinforcement-driven plasticity^18,38^ and is required for motor learning^39–41^. The BF is the source of cortically-projecting cholinergic neurons^42^, however it was unknown if BF neurons respond to VNS. To address this question, we implanted tetrodes into the BF of mice with implanted VNS cuffs (**Fig. 3a**). Extracellular activity was recorded during VNS in awake animals in their home cage (30 Hz, 0.6 mA, 100 µs pulse, 500 ms train). VNS modulated the firing rate of BF neurons, as compared to baseline firing rate (**Fig. 3b,c**). VNS altered activity in 43% of recorded units, with 61% of VNS-responsive units (28% of all units) showing increased activity relative to baseline (**Fig. 3d**). On average, the firing rate modulation of activated neurons began during the stimulation train (200 ms after stim onset) and persisted for ~1s after stimulation ended (**Fig. 3e**).

**Figure 3.**
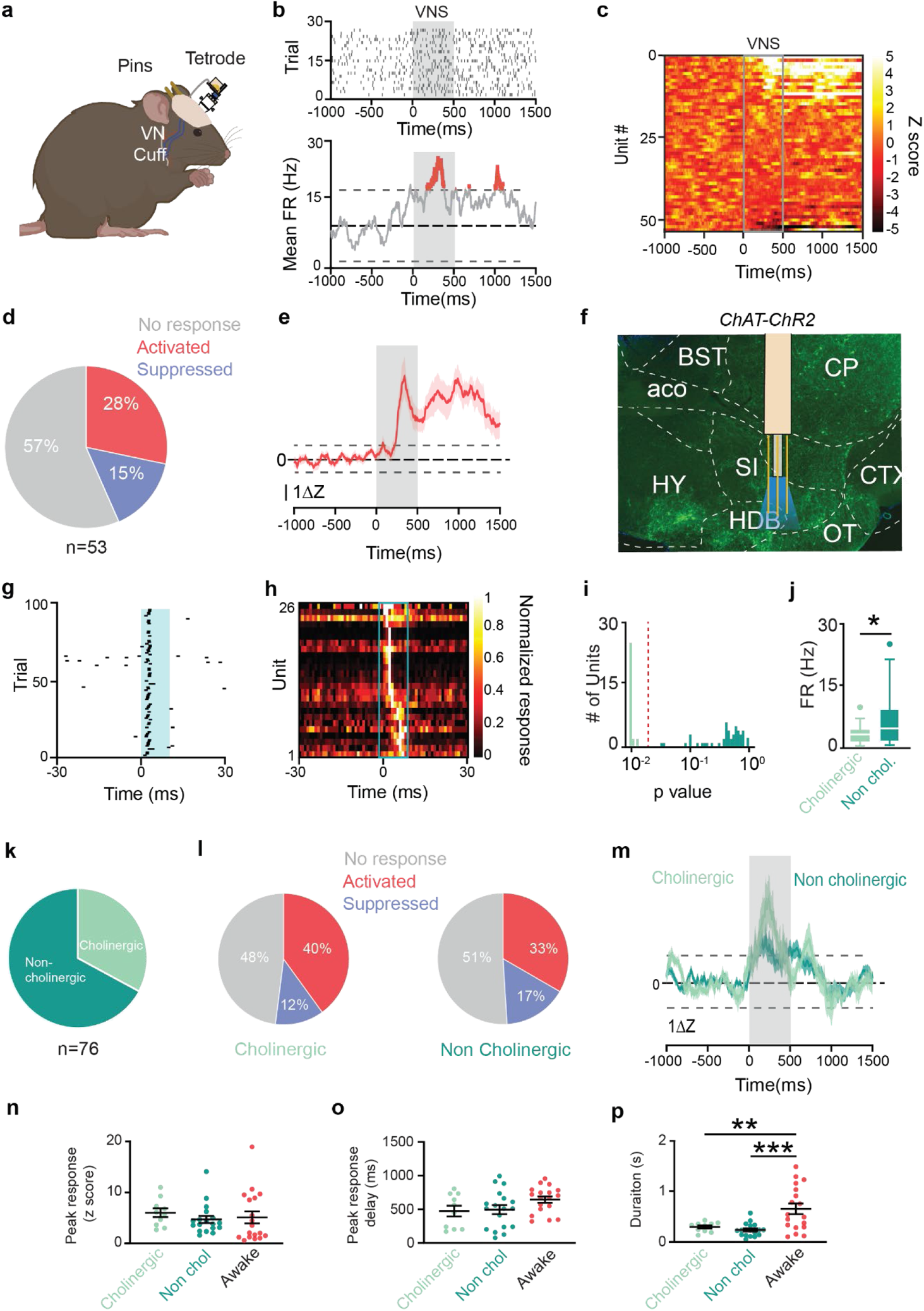
VNS drives BF neural activity in anesthetized and awake mice. **a,** Tetrodes were implanted in the left BF of mice and recordings were obtained during awake behavior. **b,** Example raster (top) and average firing rate from a response to VNS. Grey box denotes stimulus delivery. **c,** Average responses of all recorded neurons to VNS (grey box). **d,** Percent of neurons that respond to VNS (N = 5 mice, 53 neurons). **e,** Average activity of all ‘activated’ neurons in response to VNS. Dashed lines mark significance, shading represents SE. **f,** Extracellular recordings were obtained using optrodes were targeted at the HDB in *ChAT-ChR2* transgenic mice while under light anesthesia. Green fluorescence denotes the presence of ChR2. **g,** Example cholinergic neuron responding consistently to pulses of 488 nm light. **h,** Average activity of all cholinergic neurons during opto-tagging. Each row represents a neuron. **i,** Stimulus-associated latency tests (SALT) separate light responsive neurons from non-light responsive neurons. **j,** Mean baseline FR of cholinergic and non-cholinergic neurons (p=0.013, N = 5 mice, 53 neurons). **k,** Percent of neurons categorized as cholinergic (light green) and non-cholinergic (dark green) (N = 6 mice, 76 neurons). **l,** Percent of units that are VNS-responsive in cholinergic (left) and noncholinergic (right) populations. **m,** Average response to VNS for all ‘activated’ neurons. **n,** Mean peak activation during VNS. **o,** Average delay of peak activation from VNS onset. **p,** Mean duration of significantly elevated activity after VNS (cholinergic vs. awake p=0.0087, non-cholinergic vs. awake p=0.0003).

To identify cholinergic neurons from the multiple cells types found within the BF^43^, we used an opto-tagging approach^44^ combined with tetrode recordings to interrogate their response to VNS. Acute recordings were performed in anesthetized *ChAT-ChR2* transgenic mice (B6.Cg-Tg(Chat-COP4*H134R/EYFP,Slc18a3)6Gfng/J; **Fig. 3f**). Cholinergic neurons were identified by their rapid response to light (**Fig. 3g,h**) and confirmed using SALT analysis^18^ (latency 5.2 ± 1.3 ms; **Fig. 3i**). Cholinergic neurons exhibited a lower baseline firing rate (3.4 ± 2.1 Hz) than the non-cholinergic population (6.8 ± 6.2 Hz; **Fig. 3j**). Of 76 units, roughly ⅓ were cholinergic (**Fig. 3k**) and half of both neuron populations were VNS-responsive (52% of cholinergic, 49% of non-cholinergic; **Fig. 3l**). Of the VNS-responsive units, most cholinergic units and non-cholinergic units showed increased activity (**Fig. 3m**), suggesting that VNS increases activity in both cholinergic and non-cholinergic populations of the BF.

A comparison of VNS-driven activation in anesthetized recordings to awake recordings suggests that the response to VNS depends on arousal state (**Fig. 3e,m**), consistent with prior findings^14^. While the peak response magnitude and timing to VNS do not change between awake and anesthetized animals (**Fig. 3n,o**), awake animals have a longer response than anesthetized animals (awake: 652.7 ± 439 ms; cholinergic: 293.7 ± 94 ms; non-cholinergic: 235.8 ± 127 ms; **Fig. 3p**). These data confirm that VNS increases BF cholinergic activity, making it a strong candidate for mediating learning effects.

### Optogenetic cholinergic inhibition prevents VNS-enhanced motor learning

Having established that VNS can drive BF cholinergic neurons, we next wanted to determine if these neurons mediate the effects of VNS to enhance motor learning. To do so, we used optogenetic control to silence cholinergic neurons during VNS. An inhibitory opsin (AAV-EF1a-DIO-eArch3.0-EYFP) was injected into the BF of *ChAT-Cre* transgenic mice, followed by implanted optical fibers and VNS cuffs (see methods; **Fig. 4a**). Mice then learned to perform the skilled reach task (**Fig. 4b**). One cohort received Success VNS, a second received Success VNS simultaneously with optical inhibition of the BF (Arch+VNS), while the control cohort was unstimulated (**Fig. 4c**). VNS animals (40.55 ± 7.1%) performed significantly better than controls (31.23 ± 10.4%), while Arch+VNS animals performed at control levels (25.59 ± 9.3%; **Fig. 4d,e**). While all cohorts learned the task (**Fig. 4f**), cholinergic inhibition prevented VNS-driven performance increases in both learning phases (Early: 19.02 ± 9.5%; Late: 28.66 ± 4.1%) compared to VNS mice (Early: 32.81 ± 17.1%; Late: 48.28 ± 11.6%; **Fig. 4g**), demonstrating that phasic cholinergic signaling is necessary for VNS-enhanced motor learning.

**Figure 4.**
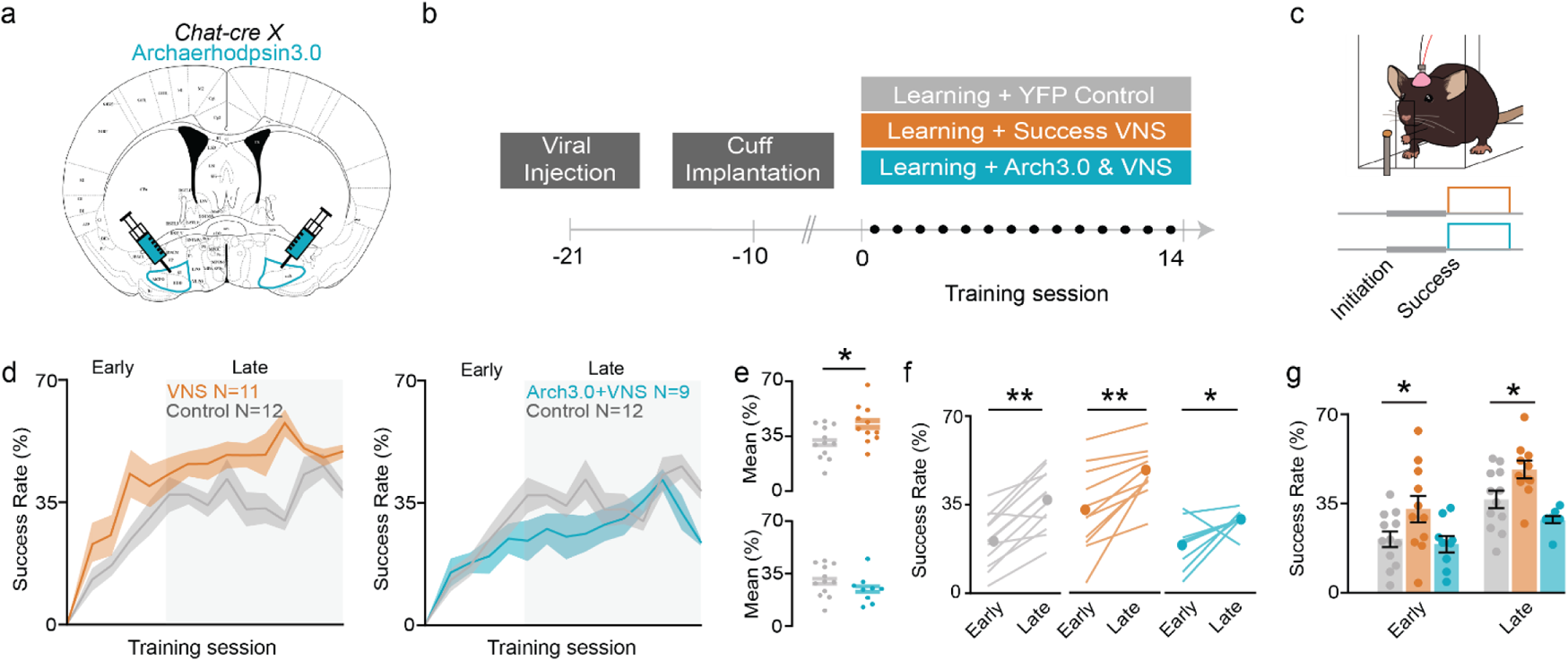
Success-paired VNS motor learning enhancement requires cholinergic neuromodulation. **a,** A subset of VNS-implanted *ChAT-Cre* transgenic mice also received injections of viral constructs containing Archaerhodpsin3.0 bilaterally in the BF (right). Mice were chronically implanted with bilateral fiberoptic cannulas for light delivery. **b,** Timeline of experimental set up and training. Each training session lasts for 20 minutes. **c,** Depending on cohort, mice receive VNS, or continuous 532 nm light and VNS simultaneously, after successful reach attempts. **d,** Average success rate for all mice over the course of learning (VNS N=11, Arch+VNS N=9, Control N=12). **e,** Mean performance across all days between VNS and control (p=0.0409 (top)), and Arch+VNS and control (p>0.05 (bottom)). **f,** Mean success rate of all groups between early and late phases (Control p=0.0001, VNS p=0.0009, Arch VNS p=0.0379). **g,** Mean success rate for control and VNS mice during early (p=0.0458) and late (p=0.0001) learning phases and control and Arch+VNS mice (p>0.05).

### VNS reduces off-target failures through increased reach consistency

To further explore how VNS can influence the learning of a skilled reach, we measured the kinematic features of the reach across learning and conditions. To obtain accurate kinematic measures, we designed a custom closed-loop automated reaching apparatus (CLARA) to track forelimb kinematics in real-time, and apply closed-loop VNS following automated classification of reach outcome^45^ (**Fig. 5a-c**). Using video data acquired by CLARA, trials were categorized into one of four categories: success; reach failures; grasp failures; and retrieval failures (**Fig. 5d**). These reach-types were compared in control, Success-VNS and Arch+VNS mice to determine if VNS influences reach kinematics and improve motor learning. Out of all errors, success-VNS mice made fewer reach failure errors than control and Arch+VNS mice (VNS: 54.14 ± 10.4%; Control: 72.16 ± 7.0%; Arch+VNS: 71.66 ± 4.5%), and more on-target grasp errors (VNS: 38.19 ± 9.2%, Arch+VNS: 24.00 ± 4.8%, control: 22.14 ± 6.0%; **Fig. 5e,f**). The reduction in off-target reach failures implies an improved accuracy in reach trajectory. Therefore, we explored if VNS drives a speed/accuracy trade-off^46^. We measured reach endpoint accuracy and outward reach velocity (see methods; **Supp. Fig. 3a,b**) and found that Success VNS does not alter endpoint accuracy or speed of reach attempts (**Supp. Fig. 3c**), suggesting that it does not influence performance through modulation of either speed or endpoint accuracy.

**Figure 5.**
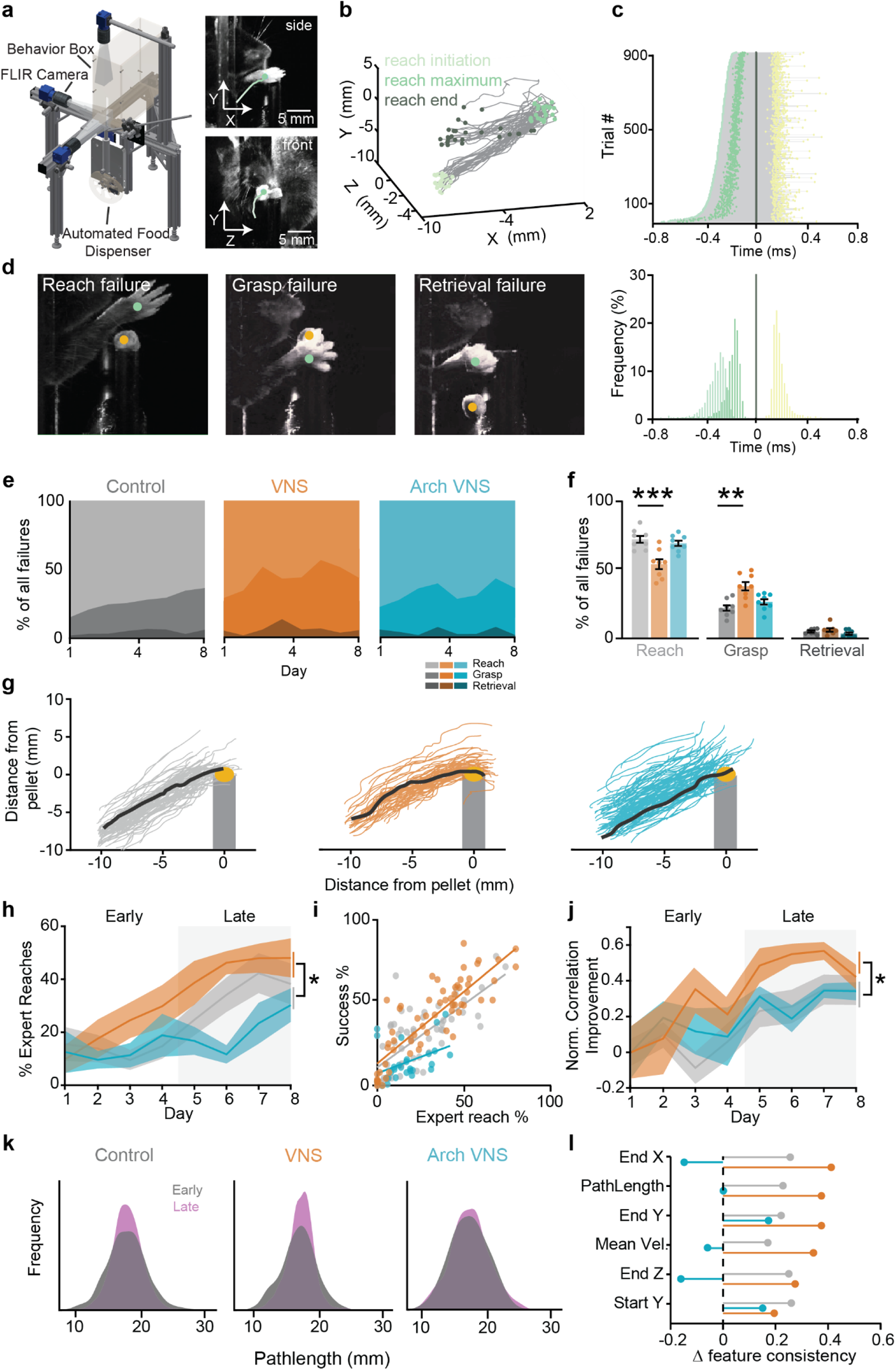
VNS improves performance through improved consolidation of reach trajectory. **a,** The Closed-loop automated reaching apparatus (CLARA) provides 3D tracking of the paw and pellet. **b**, A custom CLARA data pipeline automatically identifies reaching events (one mouse one control session). **c**, CLARA rapidly delivers stimulation after reach end in a closed-loop fashion (180±5ms). Top: duration of all stimulated control trials, yellow dot denotes stimulus delivery. Bottom: a histogram of reach timepoints normalized to reach end. **d**, Example images of the three subcategories of failed reaches (see METHODS). **e**, Breakdown of failure outcomes for each group over 8 days of learning. Light colors: reach failures; intermediate: grasp failures; dark: retrieval failures. **f**, A comparison of types of failed attempts between control and VNS (reach errors: p=0.0005; grasp errors: p=0.0035) and between control and Arch+VNS mice (p>0.05) (VNS N=8, Arch+VNS N=8, Control N=8). **g**, Examples of all outward trajectories (reach initiation – reach maximum) during a session on day 8. Black lines represent each mouse’s ‘expert reach’. **h**, Percent of reaches that are ‘expert’. Comparisons were made for the mean ‘expert’ reaches in the late learning phase (grey box) between control and VNS mice (p=0.0142) and control and Arch+VNS mice (p>0.05) (VNS N=8, Arch+VNS N=6, Control N=8). **i**, Correlation of ‘expert’ reaches and task performance for all mice, R^2^=0.62. **j**, Improvement in reach failures toward an expert trajectory as measured by increase in correlation coefficient (normalized to day 1). Comparisons were made during late learning between control and VNS mice (p=0.0455) and control and Arch+VNS mice (p>0.05). **k**, Distribution of trajectory lengths from all failure attempts during early (grey) and late (purple) learning phases. **l**, Normalized improvement in reach features from early to late learning phases.

As animals learn the skilled reach task, their reach trajectories become more similar to their final expert reach trajectory^34,47^, indicating that the animals are learning to execute a successful motor plan. To determine if VNS can influence the motor plan selection, an expert trajectory was defined for each mouse based on the average successful reach trajectory over the last two days of training (see methods; **Fig. 5g**). Across all training days, expert reaches were identified by having a >0.95 correlation with the expert trajectory. On day 1 of training, all cohorts have similar percentage of expert reaches (**Fig 5i**; **Supp. Fig. 3d**), but during late learning, VNS mice made significantly more expert reaches compared to control mice (VNS: 45.22 ± 4.4%; Control: 34.22 ± 6.6%) while cholinergic inhibition prevented this increase in expert reach selection (23.58 ± 6.2%; **Fig. 5h**). Expert reach attempts correlate strongly with behavioral performance (p=0.0001, R^2^=0.621), demonstrating that this increased stereotypy onto the expert trajectory explains the improved performance (**Fig. 5i**). VNS also shapes the trajectory of reaches that end in failure. While reach failures rarely qualify as expert reaches (**Supp. Fig. 5e**), VNS increases the correlation of reach failures to the expert trajectory during late learning to a greater degree than control mice, and cholinergic inhibition prevents this increase (VNS: 50.18 ± 16.6; Control: 29.65 ± 19.9; Arch+VNS: 33.19 ± 11.6; **Fig. 5j**). Additional kinematic features also show a VNS-driven increase of the consolidation of other reach features between early and late phases (narrower histogram with a higher peak; **Fig. 5k,l**), indicating increased stereotypy in the VNS cohort. This data suggests that VNS drives all reaches closer to the expert reach trajectory, and this is mediated by cholinergic signaling. This demonstrates that VNS improves task performance by enhancing the selection of a success motor plan.

### VNS drives acute neural suppression and activation in motor cortex

VNS paired with forelimb movement alters motor cortical map plasticity^20^, but the effect of VNS on neuronal function in motor cortex is unknown. Given that neural activity in M1 is required for both motor skill learning and execution^47,48^ (**Supp. Fig. 4a**), we hypothesize that VNS will modulate the neural activity and movement representation in M1. To investigate the effects of VNS on M1 neural activity, we imaged activity in neurons expressing the calcium indicator GCaMP6m using a head-mounted miniature microscope (UCLA miniscope V3, http://miniscope.org; **Fig. 6a**). VNS was applied to freely-moving animals in the homecage environment.

**Figure 6.**
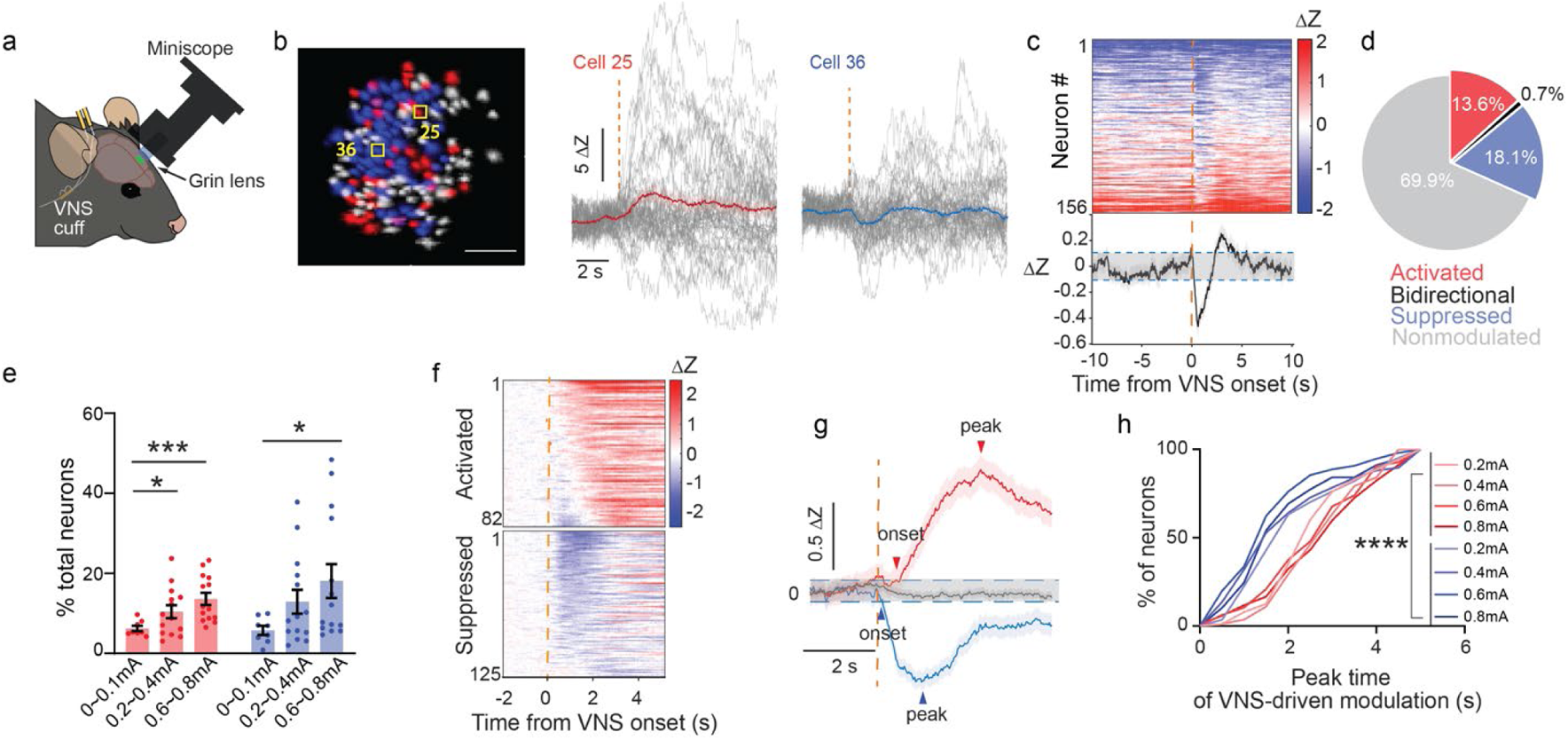
VNS drives acute neural suppression and activation in forelimb motor cortex. **a,** Placement of the grin lens and miniscope above contralateral M1, with VNS cuff in the neck and its wire connectors on top back of skull. **b,** Left, representative neural ROIs from the field of view of one mouse M1 (n = 156 neurons): neurons are pseudo color labeled as VNS-activated (red) and suppressed (blue), scale bar = 100 µm. Middle and right: two representative neurons’ Ca2+ responses aligned by VNS onset (gray: individual trials; red: VNS-activated; blue: VNS-suppressed). **c,** Top: individual neurons’ average response z-scored to inactive phases of all neurons from the representative mouse in **b**; bottom: average neural responses of all neurons from the same mouse. **d,** Average % of total neurons activated, suppressed, bidirectionally modulated and nonmodulated after 0.6 to 0.8mA VNS delivery (N = 7 mice, 767 neurons). **e,** % of total neurons that are activated or suppressed by VNS across different current amplitudes. (N = 7 mice, n= 747~807 total neurons from each stimulation group, One-way ANOVA and multiple group comparison to 0~0.1 mA group). **f,** Neural response heatmap of all activated neurons and all suppressed neurons aligned at VNS onset. **g,** Average neural activity of all activated neurons and all suppressed neurons aligned at VNS onset (N=7 mice, 82 activated neurons, 125 suppressed neurons, 0.6mA VNS). **h,** Cumulative distribution of neural response peak time of activated and suppressed neurons measured as the peak value of the average trace 0 to 5 s after VNS onset (82 to 151 neurons from each group, Kruskal-Wallis test followed by Dunn’s multiple comparisons test, p < 0.0001). *p < 0.05, **p < 0.01, ***p < 0.001, ****p < 0.0001; bars and error bars represent the mean ± s.e.m.

In response to VNS, some neurons demonstrated either activation (red, cell 25; **Fig. 6b**) or suppression (blue, cell 36; **Fig. 6b**), without a change in the overall firing rate of the neuron population (**Supp. Fig. 5a,b**). Approximately 30% of all neurons showed acute response to a VNS delivery, with roughly similar percentages of neurons showing activation and suppression (activation: 13.8 ± 5.8%; suppression: 18.1 ± 15.8%; **Fig. 6d**), and only a small fraction (0.7 ± 1.4%) showing bidirectional modulation (**Fig. 6d**). The number of neurons modulated depended on stimulation intensity (**Fig. 6e**).

Across the population of VNS-responsive neurons, we observed a temporal relationship between activation and suppression (**Fig. 6c,f**). The mean peak timing of the suppression precedes the activation by 1.2 s (Suppression: 1.6 s from VNS onset; Activation: 2.8 s; **Fig. 6g**), a relationship consistent across a range of behaviorally relevant stimulation intensities (**Fig. 6h**). Similarly, the onset of suppression preceding activation by 0.6 s (**Supp. Fig. 5c**). These together suggest that in the forelimb region of the mouse primary motor cortex, VNS first drives acute neural suppression, followed by activation in two separate subpopulation of neurons, without altering the mean population firing rate.

### Success VNS modifies the neural representation of reach outcome

We next examined the influence of Success VNS on movement representation during early learning. The neural activity in M1 was measured by miniscope imaging in freely moving mice as they learned the skilled reach task. Each reach was subdivided into a reach and an outcome phase using post hoc analysis (**Fig. 7a**). The reach phase includes the outward paw movement from reach initiation to reach max (~100 ms) and the return movement from reach max to reach end (~200 ms). Reach outcome was typically detected 350 ms after reach max and 200 ms after reach end. As anticipated^49–52^, the average population activity was significantly modulated during movement (reach or outcome phase; **Fig. 7b**). Nearly half of all neurons were movement modulated, with 13.3% of all neurons modulated during reach and 31.7% modulated during success outcome. For success outcome-modulated neurons, roughly half were activated, and half were suppressed (success-activated: 16.3 ± 6.4%; success-suppressed: 15.4 ± 10.2%; **Fig. 7c**). The outcome representation of success differs from failure, both at the level of the population average response (**Supp. Fig. 6 a,b**) and individual neural responses (**Supp. Fig. 6 c,d**).

**Figure 7.**
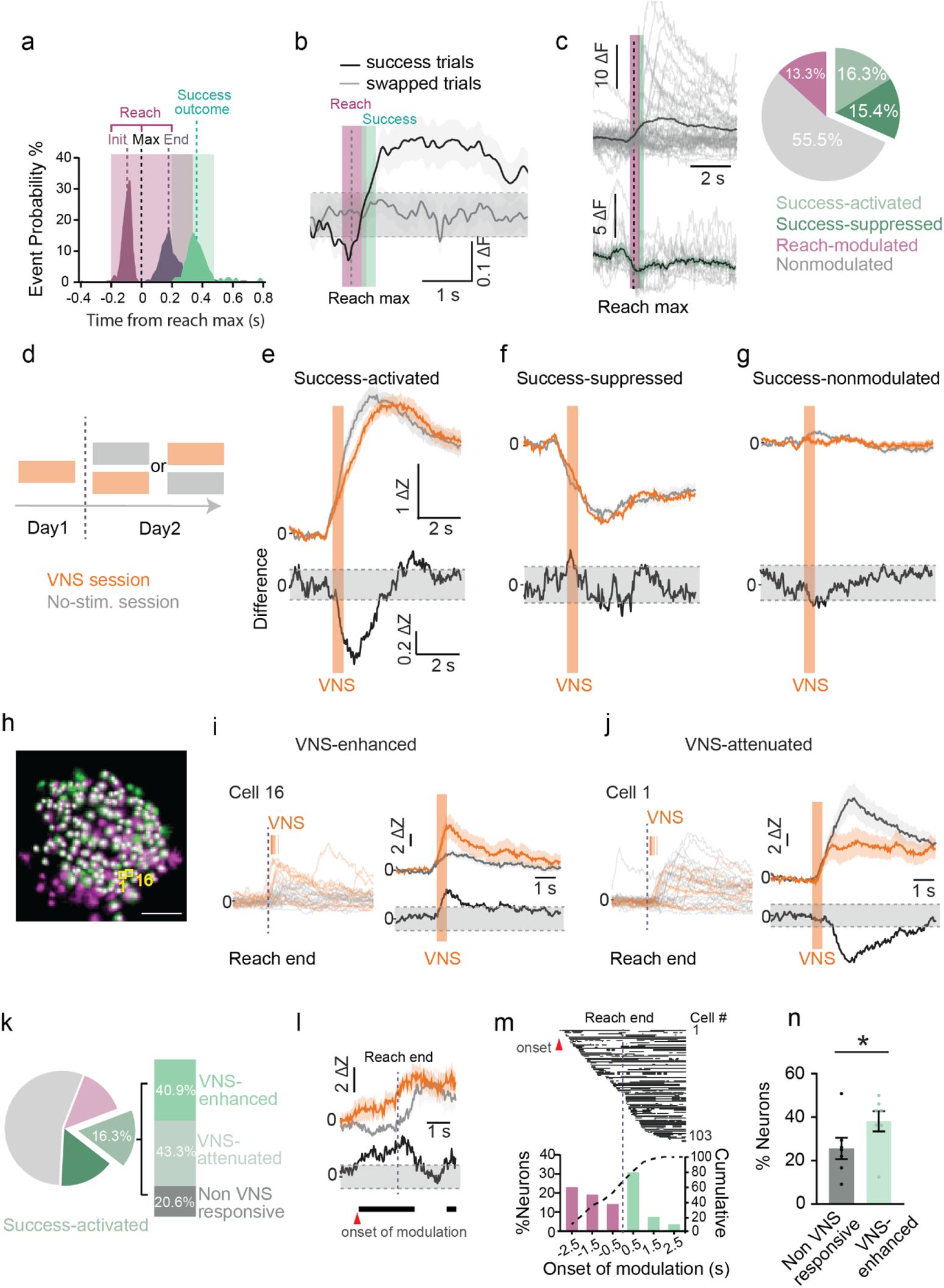
Success VNS selectively modulates activities of a subpopulation of task-activated M1 neurons in the reach task. **a**, Peri-event histogram of the task related events aligned at reach max (N = 6 control mice, day 4, n = 278 single-reach success trials). Dashed lines indicate median reach initiation −93 ms, reach max 0 s, reach end 180 ms and CLARA success recognition 360 ms. Magenta indicates full reaches, green success recognition. **b**, Average neural activity of success trials (black) and random control trials (grey, n = 488 neurons). Gray dashed line indicates 2 s.d. from the baseline mean. c, Left: representative neural responses: red success-activated, blue success-suppressed, gray individual trials. Right: % neurons modulated in the task (903 neurons; 16.3 ± 6.4% success-activated, 15.4 ± 10.2% success-suppressed, 13.3 ± 5.5% preparation/reach modulated). **d**, Assignment of VNS or no-stimulation sessions. **e-g**, Top: Average responses of success-activated (d,115 and 101 neurons), success-suppressed (e, 122 and 92 neurons) and success-nonmodulated neurons (f, 383 and 394 neurons) in VNS (orange) and in no-stimulation session (gray). Bottom: the difference trace. **h**, Registered neurons (white) in VNS (green) and no-stimulation session (Magenta). **i&j**, Left: trial responses of success-activated neurons in no-stimulation session and in VNS session (orange ticks: trials’ VNS onset). Right: the average and difference responses of the same neurons aligned by VNS onset. **k**, Percentage of success-activated neurons (n = 77 registered) that are enhanced (40.9 ± 13.7%) or attenuated (43.3 ± 21.2%) in VNS compared to no-stimulation session. **l**, One success-activated neuron modulated by VNS (Supp. Fig. 6h) also have higher neural activity before reach end in VNS session. Arrowhead: onset of increased activity. **m**, Onset timing histogram of VNS-driven modulation of success-activated neurons. n, % neurons modulated in reach in VNS-nonmodulated versus VNS-enhanced neurons. *p < 0.05, **p < 0.01, ***p < 0.001, ****p < 0.0001; bars and error bars represent the mean ± s.e.m.

During Success VNS, stimulation is delivered at reach outcome, and so the acute neural response to VNS is likely to overlap with the intrinsic response to success outcome. To accurately detect VNS-related neural activity, mice participated in two sessions of training, one with VNS and one without. These sessions were administered on a pseudo-randomized schedule (**Fig. 7d**), and the average neural response was compared between VNS and no-stimulation sessions. During VNS sessions, the success-activated neurons’ average response was first attenuated, then slightly enhanced (**Fig. 7e**). In contrast, success-suppressed (**Fig. 7f**), movement non-modulated (**Fig. 7g**) and failure-activated neural (**Supp. Fig. 6f**) responses did not differ between VNS and no-stimulation sessions. Moreover, in VNS sessions, the percentage of success activated and suppressed neurons were not significantly different from no-stimulation sessions (**Supp. Fig. 6e**), suggesting that VNS modulates neurons that already represent success outcomes. These together suggest that Success VNS specifically modulates neurons already activated by success outcome.

To track the response of individual neurons to VNS, we cross-registered neuronal ROIs between the VNS and no-stimulation sessions (**Fig. 7h**). We found that, remarkably, ~80% of the success-activated neurons were modulated by VNS (**Fig. 7k**). Despite the overall attenuation of the population response (**Fig. 7e**), nearly half of individual neuronal responses were enhanced by VNS (43.3%; **Fig. 7i,k**), while a similar portion were attenuated (40.9%; **Fig. 7j,k**). We next considered the temporal dynamics of neural enhancement and attenuation of the success outcome response, to determine if they were similar to the dynamics of activation and suppression in the home cage (**Fig. 6g,h**). Indeed, the peak attenuation occurred with similar latency to home cage VNS suppression (1.85 ± 1.20 s vs. 1.78 ± 1.10 s after VNS onset; **Supp. Fig. 6g**), followed by peak enhancement at a similar latency to peak activation in the home cage context (2.65 ± 1.50 s vs. 2.74 ± 1.20 s after VNS onset; **Supp. Fig. 6g**), demonstrating a similar temporal structure to the VNS response both during and outside of the reach behavior.

Having established that the majority of success-activated neurons are acutely modulated by VNS during success outcome, we next wanted to explore if the activity of these neurons is altered beyond the acute response to VNS. To measure neural activity across the entire reach, activity was normalized to a pre-reach baseline epoch, and movement related activity was compared between the Success VNS and no-stimulation sessions. Neural activity during the VNS session often differed across the reach phase (**Fig. 7l**), with an onset of modulation occurring prior to VNS in nearly 60% of success-activated, VNS-modulated neurons (**Fig. 7m**). More broadly, during the VNS session, all VNS-enhanced neurons are more likely to be active during the reach phase than non-VNS-modulated neurons (38.1 ± 4.7% vs. 25.6 ± 4.9%; **Fig. 7n**). Since VNS alters neural response during the entire reaching movement, this suggests that VNS effects on neural activity persist beyond those seen during acute modulation.

### VNS-driven acute neural modulation is mediated through acetylcholine receptors (AChRs)

Because the effects of VNS on motor learning are mediated by cholinergic signaling, we next set out to determine if the effects of VNS on neural activity in M1 likewise depend on acetylcholine. To test this, we injected awake, freely moving animals with a systemic acetylcholine receptor antagonist cocktail and measured the acute neural response to VNS in M1. The baseline acute VNS response in M1 was first measured in the home cage, then 15 minutes following administration of AChR antagonist cocktail, and finally in a washout session ~24 hours later (**Fig. 8a**). The percentage of VNS-modulated neurons was not changed by AChR antagonism (**Supp. Fig. 7**). However, the average response amplitude of VNS modulated neurons, both activated and suppressed, were reduced by cholinergic antagonism (**Fig. 8b-e**), demonstrating that AChR mediated signaling is required for VNS-driven acute neural activation and suppression.

**Figure 8.**
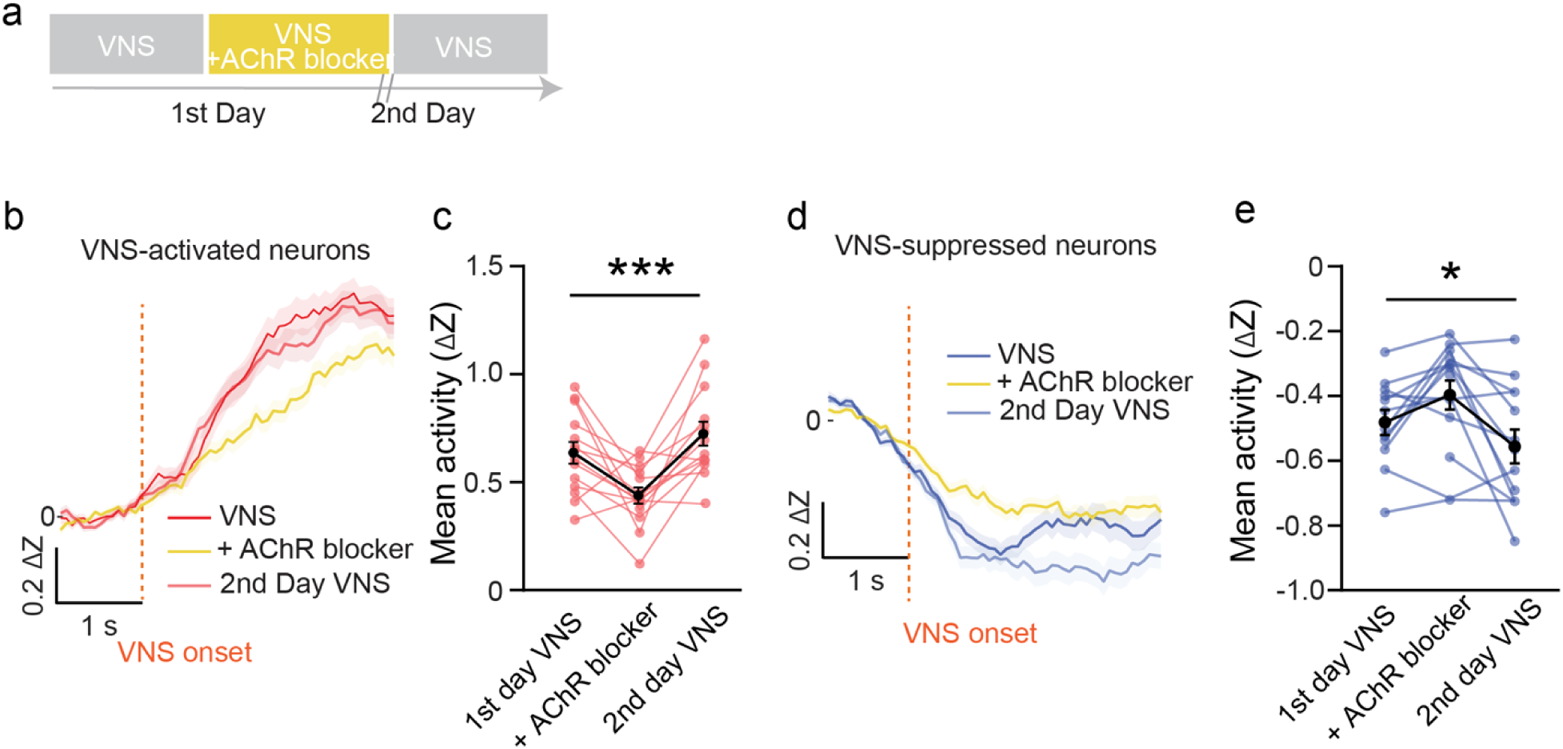
VNS driven acute neural modulation is mediated through AChRs. **a,** Diagram of experimental design: three sessions including a control VNS session in home cage, VNS session with AChR blocker, recovery VNS session on the 2nd day. **b&d,** Average neural activity of VNS-activated neurons (**b,** n = 104 ~116 neurons) or VNS-suppressed neurons (**d,** n = 124~143 neurons) in control VNS session, VNS session with AChR blocker and the 2nd day recovery VNS session. **c,** Average neural activity comparison of VNS-activated neurons quantified from 0.8 to 2.8 s after VNS onset, defined based onset and peak delay of VNS-driven neural activation in Figure. 6 (N = 7 mice x two repeats each mouse, Repeated measures ANOVA, p < 0.001). **e,** Average neural activities comparison of VNS suppressed-neurons quantified from 0.2 to 1.6 s from VNS onset defined based onset and peak delay of VNS-driven neural suppression in Figure. 6 (N = 7 mice x two repeats each mouse, Repeated measures ANOVA, p = 0.02,). *p < 0.05, **p < 0.01, ***p < 0.001, ****p < 0.0001; bars and error bars represent the mean ± s.e.m.

## Discussion

Vagus nerve stimulation paired with rehabilitation is proposed as a therapeutic treatment for a wide range of neurologic conditions, yet the mechanism by which VNS may alter neuronal activity to influence behavior remains relatively unexplored. In this study, we establish that VNS optimally enhances motor learning when paired with successful reach attempts, suggesting a reinforcement learning mechanism. Optogenetic inhibition of cholinergic neurons in the basal forebrain is sufficient to eliminate the effects of Success-VNS on motor learning and to reverse VNS-driven expert reach trajectory selection. Longitudinal *in vivo* imaging of neurons of the primary motor cortex shows that VNS selectively modulates neurons that represent reach outcome, and the effects of VNS on M1 neural activity depend on cholinergic signaling. Together, these results demonstrate that Success-VNS accelerates motor learning through reinforcement, mediated by cholinergic-dependent changes in neural representation and reach kinematics.

### VNS paired with reach success optimally enhances motor learning

To our knowledge, we are the first to demonstrate the importance of pairing VNS to movement outcome to enhance motor learning. VNS induces cortical map plasticity in healthy animals when paired with specific movements^20^ or auditory tones^21^ but has not been shown to improve task performance^20,22^. We find that VNS paired with a successful reach, but not reach initiation, enhances motor learning, indicating a role for VNS in reinforcement learning. Endogenous activity of the vagal nerve has been linked to reward and motivation^53^, and VNS in human subjects drives motivation towards reward^54^ and improves reinforcement learning^55^. However, VNS does not seem to activate the classical dopaminergic reward pathway^36^, as conditioned place preference test results indicate that VNS is not inherently rewarding or aversive. Instead, VNS may augment reinforcement cues, leading to improved selection of the expert trajectory^56,57^ and the associated neural ensembles that underlie those movements^58–60^. By augmenting reinforcement cues, VNS may help to select the appropriate neural circuits, strengthening those connections for lasting improvements in functional outcome.

The importance of VNS timing appears to differ among learning phases. During early learning, Success VNS has a unique ability to improve reach learning. Moreover, Random VNS delivered during early learning impairs reach learning, suggesting that poorly timed VNS can disrupt normal learning processes. During late learning, Success VNS maintains improved performance over control animals, while the subset of stimulated trials in Reach VNS also show performance improvements over controls (this improvement is abolished when all trials are included). In expert animals, both Success and Reach VNS provide short-term improvements over unstimulated blocks of trials. Thus, it appears critical that VNS must be paired with reach outcome during early learning, but there is more flexibility in the timing of VNS pairing during the rehearsal of a known task.

How might reinforcement-paired VNS contribute to skilled motor learning? Motor learning travels along an exploration-exploitation axis, with early exploration, expressed as motor variability, reducing as the motor behavior consolidates onto an expert solution^61–63^. Moreover, early learning motor variability predicts improved later performance of the expert motor solution^57,64,65^. This is generally referred to as an error-driven learning, in which increased exploration allows for faster identification of the expert solution. Reward is also known to influence the exploration-exploitation relationship. Generally called reinforcement-driven learning, in this process conditions of increased reward frequency lead to reduced variability and increased consolidation of movement trajectory onto an optimal motor solution^56,63,66^. Our results show that paired VNS does not increase variability in early learning, but instead improves kinematic consolidation onto an expert reach. This leads us to believe that in this context, VNS acts via reinforcement-driven learning to increase the exploitation of the expert solution, without increasing early motor variability. Future experiments are needed to explore if VNS can reinforce movements without association to a food reward.

### VNS-driven motor learning is mediated by cholinergic signaling

VNS activates multiple neuromodulatory systems in the central nervous system^16,67,68^ and the effects of VNS on cortical plasticity are mitigated with lesions of neuromodulatory nuclei, including the locus coeruleus, raphe nucleus and the cholinergic BF^15,16^. While each of these neuromodulators play a role in learning, cholinergic neuromodulatory systems are critical for use-dependent plasticity^39,69–72^, are most closely associated with reinforcement signaling^18,38^, and encode task outcome^19,73^. Lesion^17,74^ or pharmacological inhibition^75^ of cholinergic neurons is detrimental to motor learning and VNS-enhanced motor rehabilitation^9^. Leveraging the temporal precision of optogenetic suppression of cholinergic neurons, we find that a brief cholinergic inhibition is sufficient to prevent VNS-driven enhancement in motor learning. This suggests that the effects of Success VNS are mediated through phasic cholinergic signaling in the BF, and is consistent with the role of the cholinergic BF to encode outcome salience^18,19,73^.

Only limited evidence exists demonstrating an anatomical or functional connection between the vagus nerve and the cholinergic BF^76^. Using opto-tagging approaches, we were able to demonstrate robust functional connectivity between the BF and the vagus nerve. Stimulation of the vagus nerve elicited robust responses in nearly half of the cholinergic and non-cholinergic units recorded under anesthesia and more than 40% of the units recorded in awake animals. The variable timing of BF neuronal responses to VNS suggests the involvement of a multi-synaptic pathway, possibly through the locus coeruleus (LC), which is known to send direct projections to the BF^77–79^ and is activated by VNS^13,80^.

### Motor cortical neurons are modulated by VNS via cholinergic activity

Neurons in the primary motor cortex represent movement preparation and execution for dexterous movements^48,81,82^. During motor learning, these neural representations are updated to improve motor output^51,83,84^ by incorporating feedback from error and reinforcement signals generated throughout multiple regions of the central nervous system, including cortical regions^58,85–87^. Recent work demonstrated that, in addition to movement preparation and execution, M1 pyramidal neurons also report movement outcome. Neurons in superficial L2/3 of M1 represent success and failure, independent of kinematics and the food reward consumption^52^. Our data demonstrate that Success VNS attenuates the population representation of a success outcome by selectively modulating success-outcome responsive neurons. This same neural population is also more likely to have altered representation of movement preparation and reach execution, suggesting that VNS modulates neural activity beyond the acute response to stimulation. The specificity of the population of neurons that are modulated by Success VNS may indicate that VNS adds selectivity to outcome representation, which optimizes outcome signals for enhanced learning.

The relative selectivity of the effects of Success VNS on movement representation are somewhat in contrast to recent observations of widespread, long-lasting excitatory responses to VNS^14^. However, in the previous study, VNS elicited locomotion and whisking, both of which correlate to increased general arousal and widespread cortical activation^88–90^. This makes it difficult to disentangle direct VNS effects from changes in arousal. In contrast, another recent work demonstrated a VNS-driven suppression of neural response to an auditory tone that persisted even after arousal state was regressed from the neural response^91^. Our trial design controls for movement state, as the comparison between the VNS trial and no-stimulation trial occurs at the exact same reach position, and VNS did not produce any noticeable acute motor response. By eliminating the confound of behavioral states such as locomotion or quiet resting, we can detect that VNS produces an initial suppression of the outcome response followed by excitation. This effect is seen only in neurons that respond to outcome, indicating that when applied during a reach, VNS acts on a specific population of neurons that are already engaged in the representation of reach outcome.

### Optimizing VNS to treat neurological conditions

Through an improved understanding of the mechanisms of VNS, the use of this therapy to treat a range of neurological conditions can be optimized to increase clinical efficacy. For example, the recently approved VNS therapy for stroke pairs stimulation with movement, perhaps could be enhanced by pairing VNS only with movements that meet a success criterion. Similar strategies could be implemented for movement rehabilitation to treat spinal cord injury or peripheral injury. In addition, our results point to a concern that improperly paired VNS could lead to maladaptive plasticity. We find that random VNS stimulation impairs early learning. For clinical application in vulnerable populations, such as patients with neuropsychiatric conditions or in pediatric populations, maladaptive plasticity has the potential for harm. Understanding how VNS interacts with neural circuits, including both the cholinergic basal forebrain and motor cortical populations, allows for exploration of stimulation protocols to more directly target the effect of interest. For instance, models that can predict how stimulation protocols will engage specific vagal fiber types^92–94^, could be used to test differential target engagement in the brain. Alternatively, direct brain stimulation of targets, such as the basal forebrain, could be used to provide more specific neuromodulation to achieve key therapeutic results. Lastly, less invasive techniques, such as auricular VNS might still be able to convey therapeutic benefits if their stimulation protocols and target activation within the central nervous system is optimized^95,96^. The data presented here provide a framework for dissecting the role of VNS within specific therapeutic indications, with the ultimate goal of improving therapeutic delivery and patient outcomes.

## Conclusion

In conclusion, we demonstrated that VNS augments reinforcement cues to enhance skilled motor learning and accelerate kinematic consolidation on an optimal motor plan in healthy animals. VNS alters neural coding of outcome in a select neural population in motor cortex and modulates neuronal activity of this population across the entire reach. The behavioral, kinematic, and motor representation effects of VNS are mediated by phasic cholinergic activity. Understanding the behavioral and circuit mechanisms of VNS allows for future optimization of rehabilitation protocols and new avenues for the use of cholinergic manipulation to treat neurologic conditions.

## Acknowledgments

This work was supported by grant from DARPA Targeted Neuroplasticity Training (TNT): HR0011-17-2-0051. The Optogenetics and Neural Engineering (ONE) Core at the University of Colorado School of Medicine provided engineering support for this research. The IDEA Core provided data pipeline and software support for this research. The Animal Behavior Core provided support for conditioned place preference experiments. These cores are part of the NeuroTechnology Center, funded in part by the School of Medicine and by the National Institute of Neurological Disorders and Stroke of the National Institutes of Health under award number P30NS048154. We thank Sean Hansen for technical assistance in Matlab and manuscript proofreading; Eashan Sahai, Nicole Chen and Melanie Zhou for technical assistance in mice training and behavior video curation and Benjamin Temple for early technical assistance with optogenetic procedures.

## Author Contributions

SB, JH, XP and CW contributed to the text of the manuscript. JH, SB and CW designed and conducted behavioral tests. SB, JH, DD and CW designed and conducted electrophysiological recordings. XP and CW designed and conducted calcium imaging experiments. XP and CW designed and conducted pharmacological experiments. WRW and SB constructed and tested the CLARA behavioral system. KW conducted CPP tests. SB, JH, XP and RH performed surgeries for the experiments. SB, JH, XP and CW analyzed experimental data and performed statistical tests.

---

**Supplementary Figure 1.**
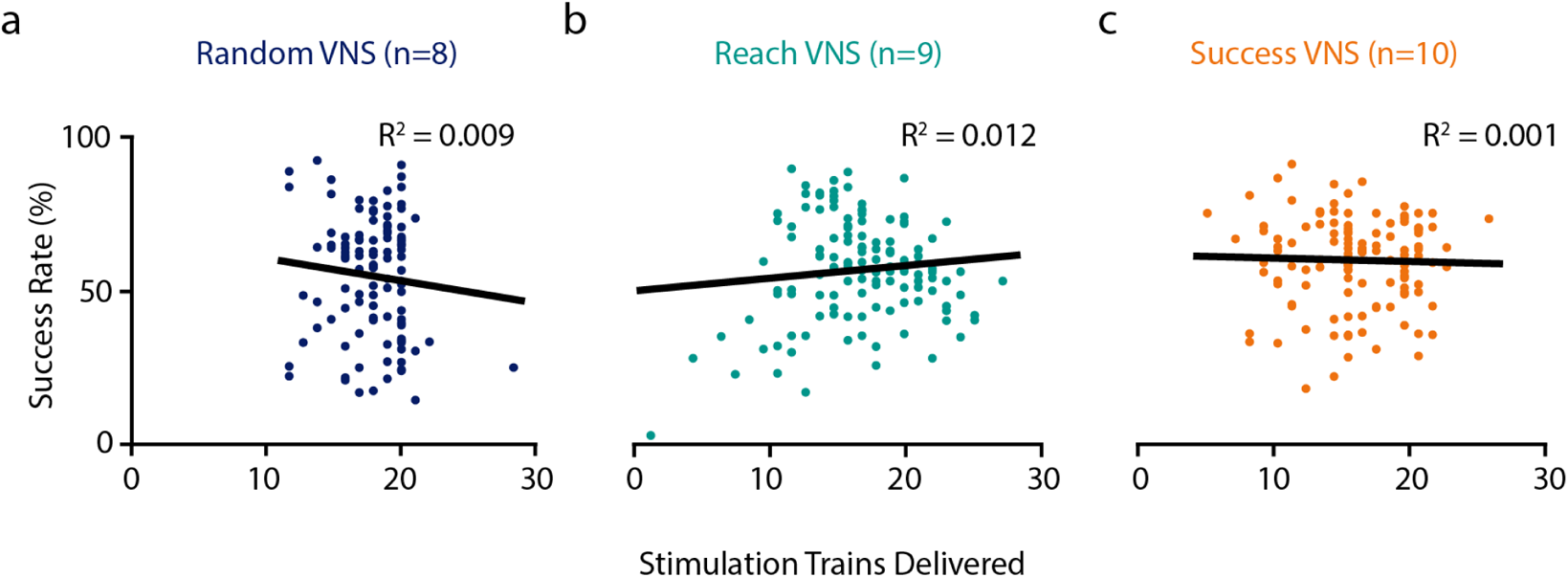
Stimulation Amount between Random VNS, Reach VNS, and Success VNS does not predict success. **a-c,** A simple linear regression was calculated to predict success rate based on the amount of stimulation trains given across groups in a particular session. **a,** Random VNS linear regression (F=1.03, p=0.3117), R^2=0.009. **b,** Reach VNS linear regression (F=2.017, p=0.22), R^2=0.011. **c,** Success VNS linear regression equation (F=0.11=, p=0.7359), R^2=0.001.

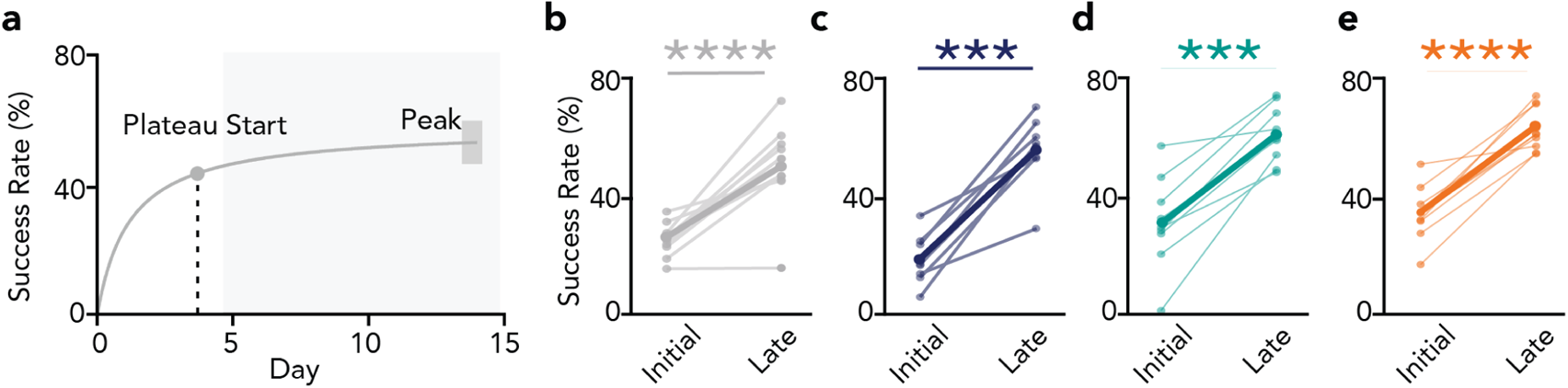
**a,** Nonlinear model of sham animal learning trajectories used to identify early and late learning phases (see methods). **b-e,** Comparison of performance during early and late learning phases**. b,** Sham VNS (paired T test, n=12, p=0.0001). **c,** Random VNS (paired T test, n=8, p=0.0001). **d,** Reach VNS (paired T test, n=9, p=0.0002). **e,** Success VNS (paired T test, n=10, p=0.001).

**Supplemental Figure 3.**
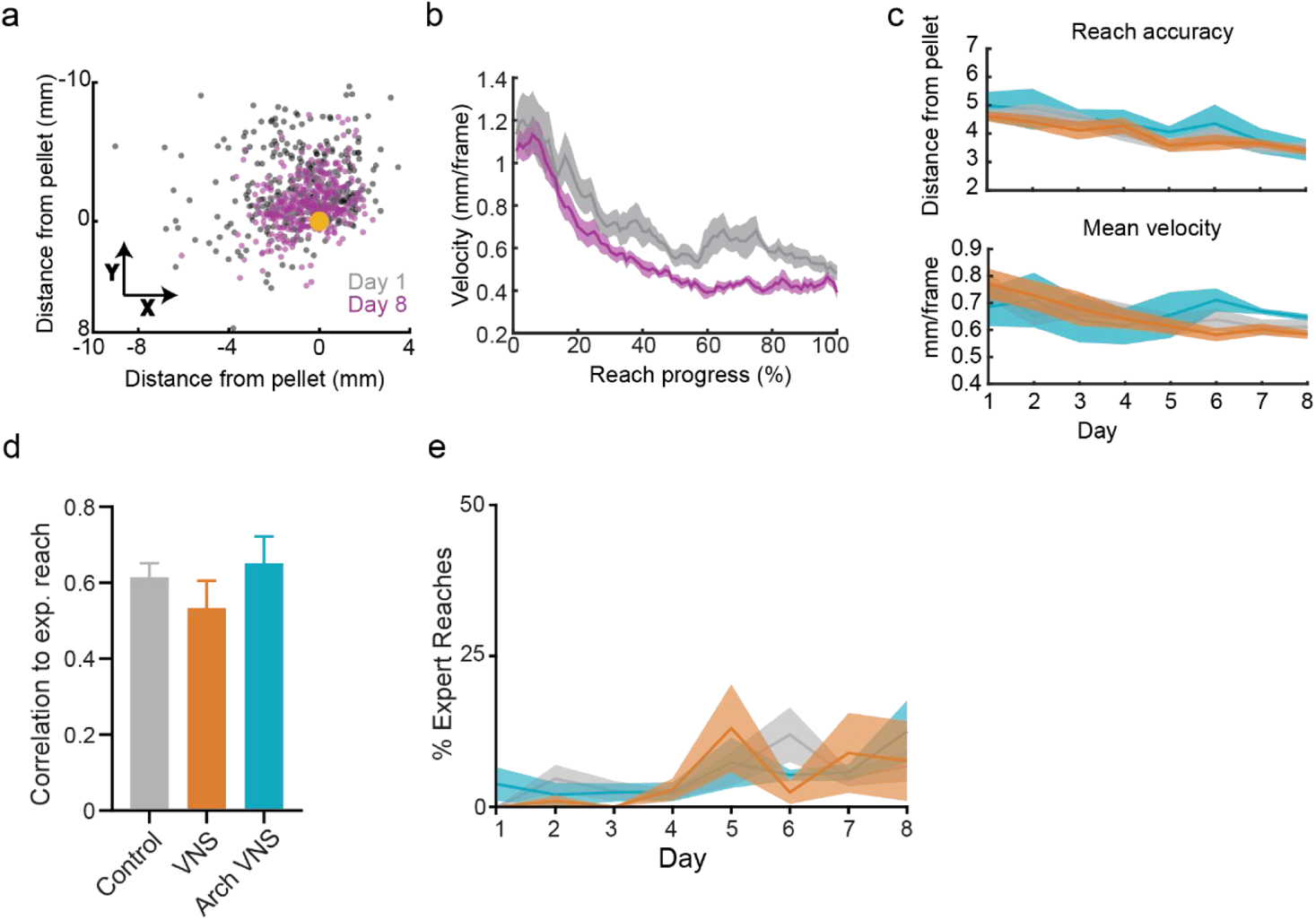
VNS does not change speed, accuracy or initial exploration. **a,** Endpoint targeting of all control reaches on day 1 (grey) and day 8 (purple) of learning. Orange dot depicts pellet center. **b**, Mean absolute velocity between reach initiation and reach maximum on day 1 and day 8 of learning. Reaches are time warped to be an equal arbitrary length. **c,** Comparison of endpoint accuracy (top) and absolute velocity (bottom) between control, VNS and Arch+VNS stimulation (p>0.05, RM ANOVA, VNS N=8, Arch+VNS N=6, Control N=8). **d,** Reach variability on day 1 for all groups, as measured by average correlation to expert trajectory (p>0.05, one-way ANOVA, VNS N=8, Arch+VNS N=6, Control N=8). **e,** Percent of ‘reach failures’ that qualify as expert reaches over learning.

**Supplementary Figure 4.**
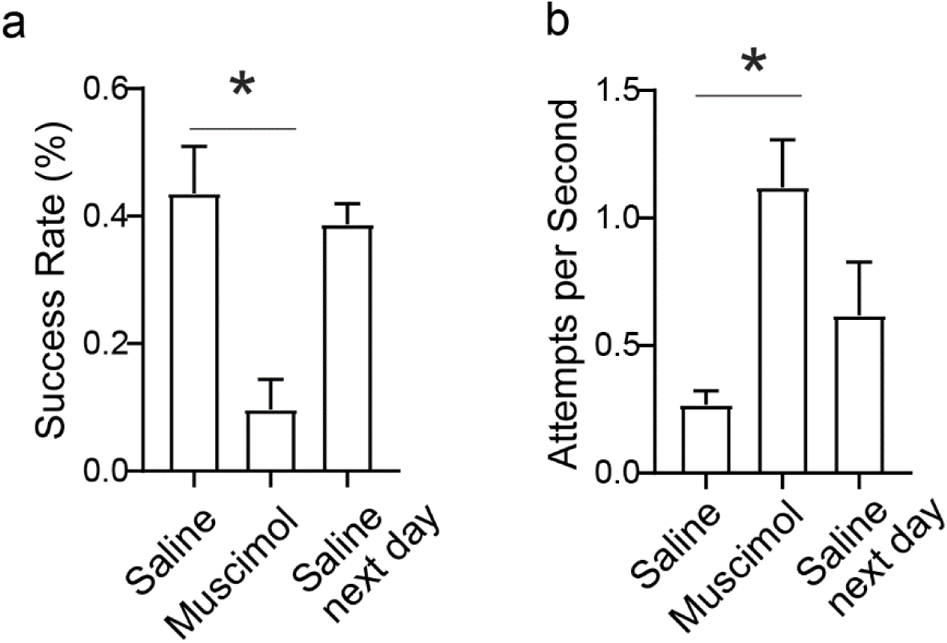
The M1 activity is necessary for successful execution of the skilled reach task. **a,** Comparison of success rate of the skilled reach task before and after muscimol infusion into M1 (N=3, one-way ANOVA, p<0.05). **b,** Comparison of reach attempts per second in the same mice before and after muscimol infusion (n=3 mice, one-way ANOVA, control ~0.3 reaches per second, muscimol 1.1 reaches, p<0.05).

**Supplementary Figure 5.**
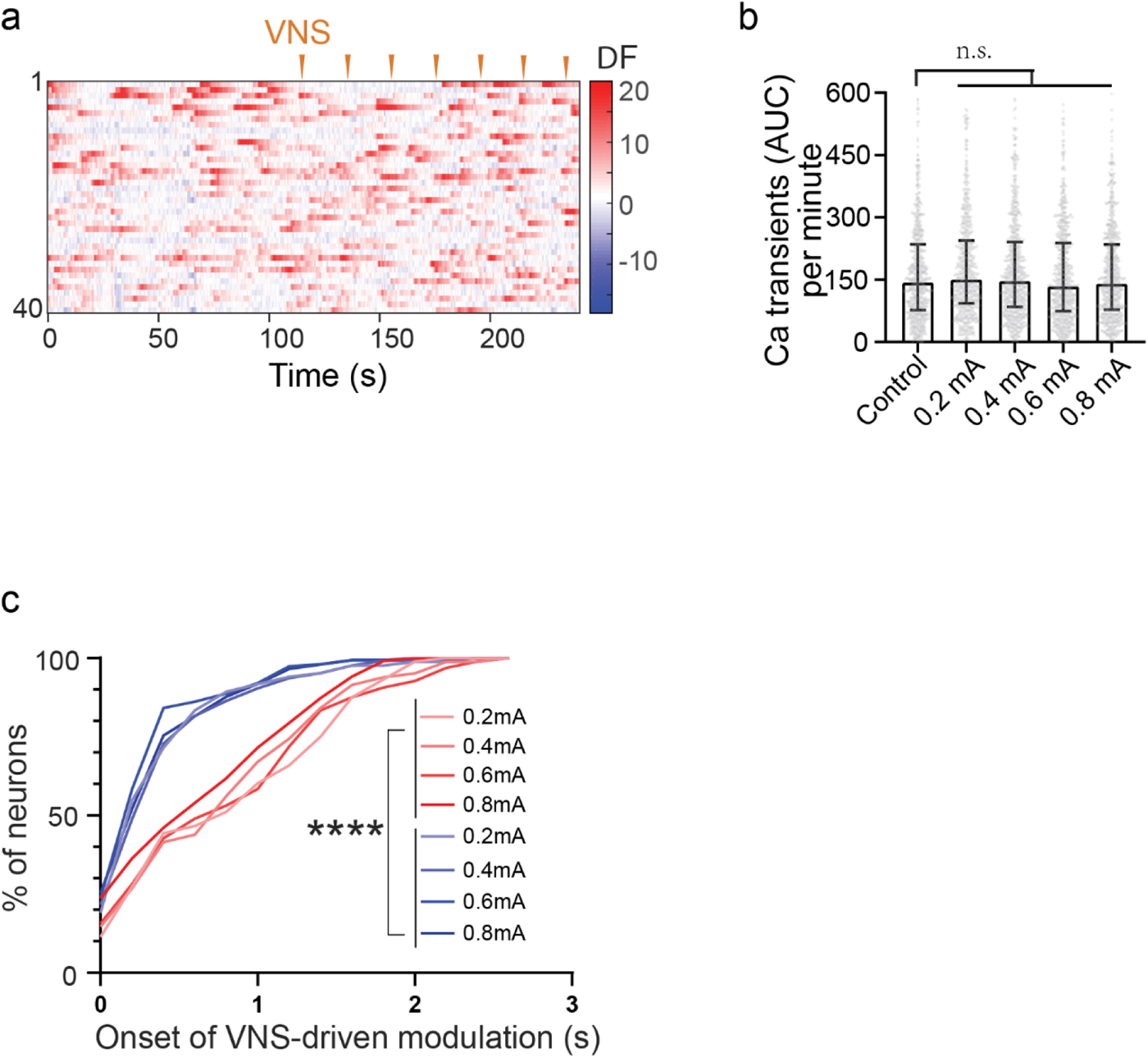
The M1 activity is constant on minutes’ time scale and the VNS response onset timing analysis. **a,** Heatmap plot of 40 neurons’ GCaMP6m calcium response in L2/3 M1 from a representative mouse in the home cage two minutes before VNS delivery and two minutes while receiving delivery of VNS trains. Each orange arrow indicates one VNS pulse train delivery (15 pulses at 0.1 ms duration, 30 Hz, 0.4 mA). **b,** Ca transients quantified as area under the curve (AUC) of the fluorescence trace per minute in the 2nd minute after VNS application starts in VNS mice (N=7 mice, 676 to 732 neurons, plot as median with interquartile range. Kruskal-Wallis test followed by Dunn’s multiple comparison, p>0.05 for all VNS stimulation groups compared to control. Controls are the same mice without VNS delivery). **c,** Cumulative distribution of neural response onset (measured as the first value goes above 2 s.d. of the baseline mean in the average trace 0~5 s after VNS onset) of VNS-activated or suppressed neurons (n=82 to 151 neurons from each group, N=7 mice, Kruskal-Wallis test followed by Dunn’s multiple comparisons test, p<0.0001).

**Supplementary Figure 6.**
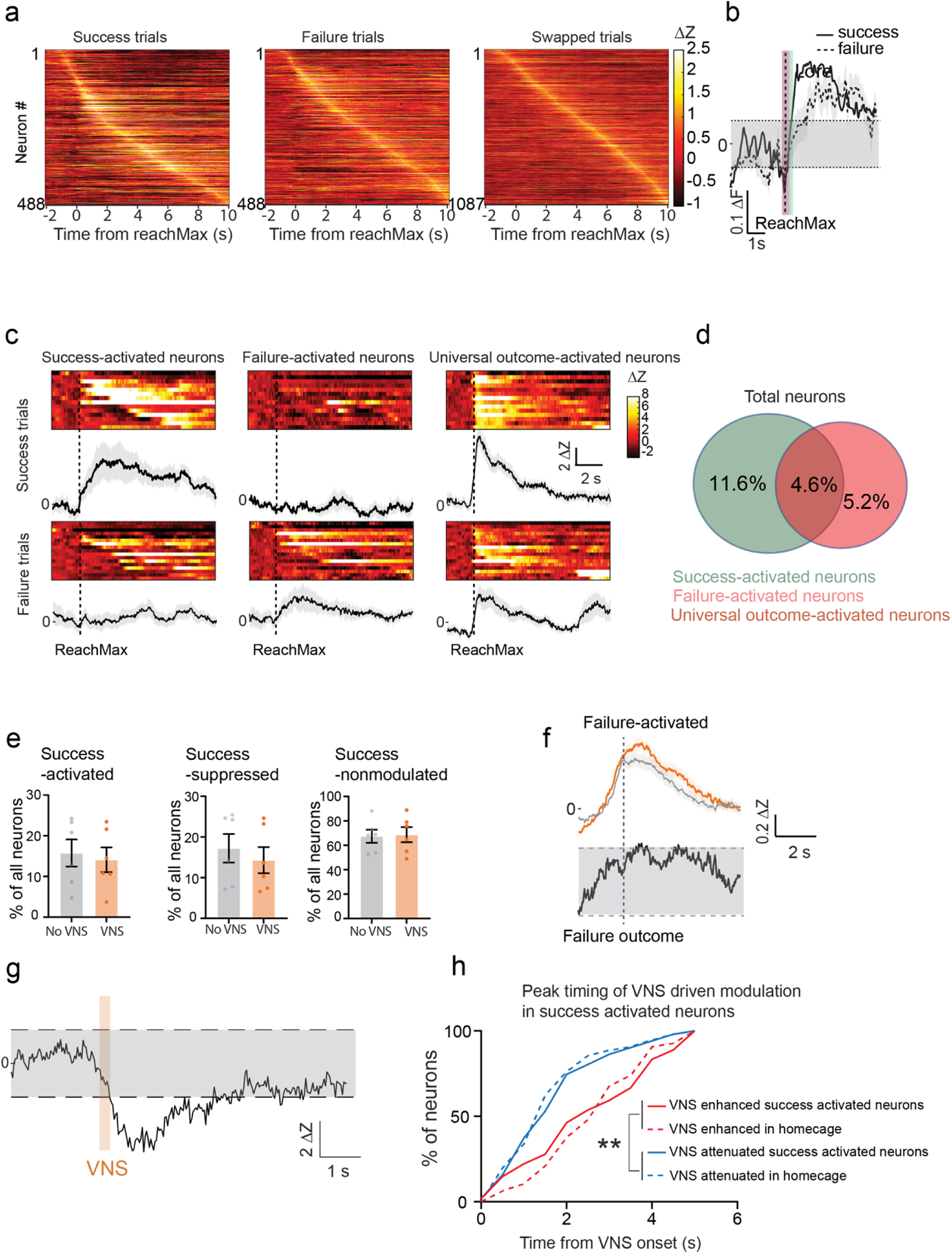
The M1 neural dynamics in the reach task. **a,** Heatmap plot of individual neurons’ response sorted by the peak timing; reach max is 0 s (N=6 mice, n=488 neurons). **b,** Average neural activity of all neurons in single-reach success trials (black line), and single-reach failure trials (dotted line). Grey block indicates 2 s.d. from the baseline mean during −6 to −3 s before reach max. **c,** Heatmap plot of trials and average response of representative success-activated neuron, failure-activated neuron and universal outcome-activated neuron aligned by reach max. **d,** % of success activated-neurons, failure-activated neurons and universal outcome-activated neurons (N=1 mouse, n=172 neurons). **e,** The percentage of success-activated, -suppressed and non-modulated neurons in VNS sessions and in no-stimulation sessions (N=6 mice, n=488 neurons). **f,** Average neural activity of failure-activated neurons in VNS session (orange, n=77 neurons) and in no-stimulation session (grey, n=73 neurons), aligned at outcome recognition by CLARA. **g,** Cumulative distribution of neural response peak time of VNS enhancement or attenuation of outcome-activated neurons’ outcome response. VNS driven enhancement or attenuation are measured as the peak value of the difference trace (individual neuron’s VNS session average – no-stimulation session average) 0~5 s after VNS onset (51~54 neurons from each group, 6 mice, Kruskal-Wallis test, p<0.0001). **h,** The response to VNS of representative success-activated neuron in fig. 7l aligned at VNS and normalized to the −3 ~0 s before VNS.

**Supplementary Figure 7.**
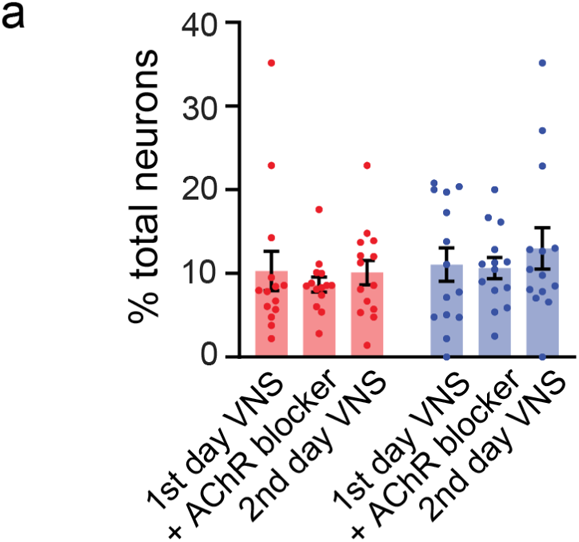
The neural populations analysis with AChR blocker. **a,** Percentage of VNS-activated neurons and VNS-suppressed neurons in the total neuron populations after 0.4~0.6 mA VNS delivery in control VNS session, VNS session with AChR blocker and recovery VNS session 2nd day (N=7 mice. n=627 neurons, Brown-Forsythe ANOVA test, p>0.5)

## Methods

### Animal care

All animal procedures and experiments were performed in accordance with protocols approved by the Institutional Animal Care and Use Committee at the University of Colorado Anschutz Medical Campus. Male and female adult C57BL/6 wild-type mice between the age of 2 and 10 months old were used for all experiments unless otherwise noted. Mice were group-housed before surgery and single-housed following surgery and throughout behavior training. Mice were kept on 14 hr light/10 hr dark schedule with ad libitum access to food and water with exception from behavior-related food restriction (Forelimb Reach training).

### Surgery

#### Vagus nerve stimulating cuff implantations

Commercial cuffs from Micro-leads (150um cuffs) and Cortec (100 um microsling cuffs), soldered to gold pins or Plastics1 connectors, were implanted on the cervical vagus nerve^28^. Mice were anesthetized with 4.5% isoflurane anesthesia for induction and maintained with 1.5%. 1% injectable lidocaine was used locally at incision sites. Eye ointment was applied to the eye to prevent corneal drying. Temperature was regulated at 37°C with a thermostat-controlled heating pad. The vagus nerve was accessed with an incision in the ventral cervical region, and the nerve was bluntly dissected from the carotid sheath. The cuff was tunneled subcutaneously to the ventral cervical incision from an incision at the base of the dorsal skull. The vagus nerve was placed in the cuff. The ventral cervical incision was sutured using 6-0 absorbable sutures. The dorsal skull was cleaned using saline and ethanol, and electrical connectors were fixed to the skull using dental cement (C&B Metabond). GLUture (WPI) was used to seal the skin around the dental cemented headcap. Stimulation efficacy was measured using peripheral biomarkers such as breathing rate changes and heart rate reduction^28^ on the day of surgery and weekly until the end of experiments with a paw sensor (Mouse Stat Jr., Kent Scientific). Mice received sub-cutaneous lactated ringers (~100 µL as necessary), intramuscular gentamicin (3 mg/kg), and intraperitoneal meloxicam (5 mg/kg) following surgery and as needed in cases of dehydration, infection, or pain. Mice were monitored for 7days to ensure proper recovery from surgery before any subsequent experiments were conducted.

#### Viral injections and optical fiber implantation

All surgeries were performed on mice expressing Cre recombinase driven by the ChAT promoter (ChAT-IRES-Cre, Jackson Labs stock #0064100^97^). Animals were prepared for surgery as described above, and the hair was removed prior to an incision over the dorsal skull. 200 µm diameter fiber optics with 1 cm ceramic cannulas were fabricated in-house using ONECore facilities. The skull at the dorsal incision was cleaned using sterile saline and ethanol. Two craniotomies were opened above the basal forebrain in each hemisphere (0.35 mm posterior, ±1.6 mm lateral of bregma) using a dental drill. Glass pipettes containing a floxed inhibitory archaerhodopsin (AAV-EF1a-DIO-eArch3.0-EYFP, UNC) or yellow fluorescent protein (YFP) control (AAV-EF1a-DIO-EYFP-WPRE-pA, UNC) were then inserted bilaterally into the basal forebrain using a stereotaxic device (−4.75 mm from dorsal surface of the brain). Approx. 210 nL of viral construct was injected over 5 minutes. The pipettes were removed and 200 µm fiber optic cannulas were inserted above each injection site (−4.65 mm from the dorsal surface of the brain). The craniotomies were then sealed using a surgical silicone (Kwik-Sil). The cannulas were fixed to the skull using dental cement (C&B Metabond). GLUture was used to seal the skin around the headcap. Mice received intramuscular gentamicin (3 mg/kg), and intraperitoneal meloxicam (5 mg/kg) following surgery and as needed in cases of infection, or pain. Mice were monitored for 7 days to ensure proper recovery. After 14 days, mice underwent VNS implantation described above (see *vagus nerve stimulating cuff implantations*).

#### Electrode implantation

##### Chronic tetrodes

Custom twisted wire tetrodes, built in-house, were implanted into the left basal forebrain of mice. Surgical preparation was as described above. Two craniotomies were opened using a dental drill: one above the left basal forebrain (0.35 mm posterior, 1.6 mm lateral of bregma) and one above the cerebellum (~1 mm posterior of lambda, midline). A tetrode was then inserted into the basal forebrain craniotomy using a stereotaxic device (−4.75 mm from the dorsal surface of the brain). A gold ground pin was inserted into the second craniotomy. Both craniotomies were sealed with surgical silicone (Kwik-Sil) and the tetrode was fixed to the skull using dental cement (C&B Metabond). GLUture was used to seal the skin around the dental cemented headcap. Mice received intramuscular gentamicin (3 mg/kg), and intra-peritoneal meloxicam (5 mg/kg) following surgery and as needed in cases of infection, or pain. Mice were monitored for 7 days to ensure proper recovery after which they underwent VNS implantation described above (see *vagus nerve stimulating cuff implantations*).

##### Acute Optrodes

Vagus nerve cuffs were implanted in transgenic mice expressing channelrhodopsin2 in cholinergic neurons (ChAT-ChR2^98^) using the protocol described above (see *vagus nerve stimulating cuff implantations*). After cuff implantation, while mice were still under anesthesia (1.5% isoflurane), mice were moved to a stereotaxic apparatus and a cranial window (2.5×2.5 mm) was opened above the left BF (0.35 mm posterior, 1.6 mm lateral of bregma). The stereotaxic apparatus was placed in a Faraday cage and single shank Optrodes from Neuronexus (A1x32-Edge-10mm-20-177-OA32LP) were inserted into the BF (−4.75 mm from the dorsal surface of the brain). Extracellular recordings were then performed while mice then underwent VNS (see *BF response to VNS*) and opto-tagging (see *opto-tagging*) protocols. Mice were sacrificed at the end of the experiment and the cuff and optrode were recovered.

#### Cranial window surgery for miniscope objective lens and baseplate installation

Mice were anesthetized with isoflurane and maintained similarly as described above until skull was exposed. A round cranial window (~1.8 mm diameter, ML 1.5 mm, AP 0.3 mm for center) was made above M1 contralateral to the reaching paw using a dental drill. A viral vector (AAV1.Syn.GCaMP6m.WPRE.SV40) was infused at 2×10^12^ titer to 200 ~300 µm beneath brain surface at 3~4 sites in the cranial window around the center, with ~200 nL at each site. Objective grin lenses (Edmund Optics, #64520, 1.8 mm, 0.23 mm WD) were lowered through the cranial window and pressed against the brain surface. The lens’ side was sealed by surgical silicone (Kwik-Sil) and secured by dental cement. The exposed part of the lens above the skull was further coated with black nail polish. 3~4 weeks later, a metal baseplate was mounted to the skull over the lens with Loctite glue (Loctite 454 prism), guided by a miniscope for optimal field of view while the mice were anesthetized with isoflurane (1.5%). After the baseplate was securely mounted, the miniscope was taken off, a cap was attached to the baseplate and the mouse was returned to the home cage.

#### Muscimol cannula surgery and infusion

Using the same craniotomy surgery procedure described above, a round cranial window (~1.8 mm diameter, ML 1.5 mm, AP 0.3 mm for center) was made above the contralateral M1 and a plastic cannula (2~3 mm long pipet tip) was inserted into the window right above the brain surface. Then the pipet tip was secured by Kwik-Sil and dental cement. The top of the cannula was sealed with Kwik-Sil if the mouse was not undergoing behavior tests within a couple of hours. Before behavior tests, the mice were briefly anesthetized with isofluorane, the seal was removed and 1~2 μL of muscimol at a concentration of 1 mg/mL was infused into the pipet tip.

### Behavior

#### Manual training of skilled forelimb reach task

Mice were trained and scored on a skilled reach task^27^. Mice were food restricted and maintained at 85-90% of their free feeding weight throughout training. Following food restriction, mice were habituated to the training box for 20 minutes where mice were given 20 mg food pellets (BIO-serv) near the window of the box where reaching occurs. The training box is a custom-built plexiglass box with a 1 cm wide opening that provides access to a post with a divot to hold a pellet located approximately 1 cm away, with a left offset from the center of the opening to force right forepaw reaching. Learning sessions then occurred for 14 consecutive days where mice perform a reach to grasp task with the right forepaw for food pellets. Rehearsal sessions occurred 7-10 days after training and featured stimulated and unstimulated trials. For both training and rehearsal sessions, mice were scored on a per trial basis until 20 successful attempts or 20 minutes passed. A trial terminates in a success, or the pellet being knocked off the pellet holder by the mouse. Trial outcomes were recorded by the trainer in real time. A success was defined by when the mouse grabbed the pellet and returned it into the cage. Errors were subcategorized into: “reach error” (failure to correctly target), “grasp error” (failure to grasp the pellet), and “retrieval error” (successful grasp of pellet, but failure to return it into the box).

### VNS experiment groups

Experiment groups for training were based on stimulation protocols. Sham VNS cohort were implanted with a cuff but were not stimulated at any point during learning. Success VNS were manually stimulated following every successful trial -- days 7 and 14 featured designated blocks without stimulation to track baseline learning levels versus trial-to-trial performance. Random VNS received stimulation at random intervals as generated by an Arduino board to achieve between 15-20 stimulations per day, matching stimulation rates to other groups. Reach VNS received manual stimulation prior to movement onset on a pseudo-random 50% subset of trials to normalize VNS trains delivered. All groups were trained on the forelimb reach task for 14 days. For performance measurements in animals trained without VNS (**Fig. 2 k-n**), Reach VNS and Success VNS were applied during daily behavioral sessions. All animals received both Reach and Success VNS sessions on different days, with randomized order of session assignments across animals.

#### CLARA skilled reach training and behavior data acquisition

Mice were food restricted and habituated identically to manually trained animals. Dimensions of the behavior box were also identical in manual and CLARA cohorts. Behavior box used for miniscope recording was modified so that the front panel had an alcove above the height of the mouse head to accommodate the miniscope when the mouse was close to the slit to reach. On day 1, the mice were primed to have one success before CLARA training session started. Learning sessions then occurred for 14 consecutive days (or specified otherwise in results), where mice perform a reach to grasp task with the right forepaw for food pellets. Each trial started as the automated dispenser placing a food pellet on the post, as the mouse reached to successfully retrieve it or knocked it off, the CLARA would mark success or failure as the trial outcome as the end of this trial. Each session lasted for 20 minutes, and mice were scored on a per attempt basis. Using the CLARA behavior system, high speed (150 Hz) video data was recorded from three FLIR Blackfly® S (model BFS-U3-16S2M-CS, Edmund Optics) cameras placed in front of the box, lateral to the box, and at a 45° angle above the box from the opposite side from the lateral camera (**Fig. 6a**). A neural network was trained prior to experiment sessions using manually annotated frames of the skilled reach behavior labeling the hand center and the pellet. Video frames from all cameras were sent through this network in real time to identify the location and state of the hand and pellet. This information was used to initiate trials via pellet placement, and to categorize attempt outcomes as either success or failure so that stimulation could be delivered in a closed-loop manner (for additional details, see^45^). The timing of pellet placement, success or failure outcome, VNS delivery, and optogenetic light delivery was recorded through CLARA. In the miniscope cohort, the timing of miniscope neural recording was cross registered with behavior video frames through a CLARA-controlled Arduino board.

#### Optogenetic+VNS experiment groups

Experiment groups for training were based on stimulation type. The control cohort was injected with a floxed YFP construct that did not contain an opsin and received light stimulation (see *light stimulation parameters*), the Success VNS cohort was electrically stimulated (see *VNS stimulation parameters*), and the Arch+VNS cohort received both light and electrical stimulation. All groups received stimulation following every successful trial automatically through the CLARA system.

#### Miniscope groups

Mice wore miniscopes for 5~10 minutes a few times in their home cages or the CLARA training box to habituate the weight. When recording VNS response in home cage, mice wearing miniscopes and VNS wires were put in home cage. About 4 minutes spontaneous neural activity were recorded as mice freely moved in the home cage, then 30~40 VNS were delivered every 20~30 s. Afterward, another ~4 minutes spontaneous activity was recorded. In sessions with AChR antagonists, scopolamine (1 mg/kg body weight) and mecamylamine (10 mg/kg body weight) were dissolved in saline and delivered to mice through intraperitoneal (IP) injection. The concentration of scopolamine and mecamylamine cocktail was chosen to have effects in brain circuits related to memory and learning without debilitating effects, according to previous studies^99,100^. 15 minutes after cocktail administration, the neural response was recorded during a home cage session.

For reach training recordings, food restricted mice were mounted with a miniscope and VNS wires and put in the training box to start a CLARA training session. The minisope acquisition was turned on immediately as the CLARA training session started and each frame of the video was cross registered with the CLARA video frames. All mice were primed without VNS to have one success reach before the first session started. On the first day, VNS mice participated in one 20-minute Success VNS session. From day 2 to day 4, each VNS mouse participated in two sessions of training, with one of them being a Success VNS session and the other a no-stimulation session in which VNS was not delivered, which was given on a pseudorandomized schedule (**Fig. 7d**). Control mice also received two training sessions without VNS each day. On days when mice receive two sessions of training, the two sessions are 1~3 hours apart.

#### Place preference test

We used a standard conditioned place preference test to examine if VNS is rewarding or aversive. The behavior apparatus contained two compartments separated by a gate. Mice were tested for baseline preference in an initial 20-minute session where mice can freely navigate between compartments. Mice then were trained for three days with two 20-minute sessions each day where they received stimulation in only one compartment. Stimulation was delivered pseudo-randomly approximately once per minute. On the day of testing, no stimulation was given, and mice were allowed to freely navigate between compartments to see which compartment they spent more time in^101^. The amount of time spent in each compartment was compared between the baseline and testing day. Experiments were conducted with assistance from CU Anschutz Behavior Core.

#### Stimulation parameters

##### VNS stimulation parameters

For all VNS experiment groups, VNS was delivered as a 500 ms train of 15 pulses, with 100 µs phase duration at 30 Hz. Current amplitudes were 0.4-0.6 mA. Stimulation parameters were controlled and delivered using Master8, PulsePal, or a CLARA+Arduino system, which were connected to a stimulation isolation unit (A-M Systems, Model 2200 Analog Stimulus Isolator) to control amperage.

##### Light inhibition parameters

For all light stimulated groups, 561 nm light was delivered continuously for 500 ms. Light was delivered through a 200 nm fiber-optic cable from a Class IIIb diode pumped solid-state laser (Cobalt) at 0.5 mW (calculated based on output efficiency from the bottom of the optical fiber). Stimulation parameters were controlled and delivered using a PulsePal, or a CLARA+Arduino system connected directly to the laser.

### Behavior and kinematic analysis

#### Manual behavior analysis

A success percentage was generated for each session of each animal by determining the number of trials that resulted in a successful retrieval out of all trials initiated. Success percentages were compared between stimulation groups across all days of training, as well as by early and late learning phases. Early learning phase refers to days 1-4 of training, while the late phase refers to days 5-14, which were defined using a Weibull growth curve (See *quantification and statistical analysis*). On days where animals received blocks of stimulation, such as rehearsal groups, stimulated and unstimulated trials were compared on a per mouse basis within days. To determine a trial-level effect for the Success VNS group, we divided all trials to three categories, pre-success trials that occur immediately before each success trial, success trials and post-success trials that occur immediately after each success trial. We then compared the success rate between pre-success and post-success trials.

#### Behavior curator analysis

Videos acquired during CLARA training sessions were processed by custom Python scripts overnight to extract key reach timepoints: reach initiation (ReachInit), when the hand leaves the box; reach max (ReachMax), the outward point of maximum distance from reach initiation; reach end (ReachEnd), when the hand returns to the cage; and stimulation onset (stim), when a trial received a trigger pulse for VNS or light stimulation. The stamps of reachInit, reachMax and reachEnd were further manually screened for consistency. The accuracy of CLARA trial outcome classifications were verified, and failures were subcategorized *post-hoc* into reach, grasp and retrieval failures (see *Manual training of skilled forelimb reach task* for failure definitions).

#### Kinematic analysis

3D location of the center of the paw and pellet were tracked during reach attempts (between reach initiation and reach end). Tracking data was extracted using custom MATLAB scripts (MATLAB Simulink) and documented as 3D data arrays for kinematic analysis. Positional data for gross targeting analysis was determined by selecting the 3D location of the hand and pellet at the reach max timepoint. Points were normalized such that the pellet center was 0,0,0. Euclidean distance between the hand center and pellet center was then calculated using the norm function from MATLAB. The mean distance from the pellet was compared in early and late phases and between stimulation groups. Reach velocity was obtained by measuring the absolute velocity between reach initiation and reach end, and then averaging the velocity over that period. The mean velocity was compared in early and late phases of training and between stimulation groups.

#### Reach consistency and expert reach

Positional data between reachInit and reachEnd were normalized such that the pellet center was 0,0,0. Reach trajectory was defined as the time between reach initiation and reach end. Each trajectory was then temporally warped to be the same arbitrary ‘length’ of time using dynamic time warping^102^. An expert trajectory was constructed for each mouse by averaging the trajectories of all successful reach attempts made on each mouse’s last two days of training. Reach consistency was determined through comparison of reach trajectories to each mouse’s expert trajectory. Reach trajectories were compared to the expert trajectory through a correlation coefficient to obtain the mean correlation coefficient for each mouse in each training session. Additionally, any individual reach that had a correlation of 0.95 or higher with the expert trajectory was defined as an ‘expert reach’, and the percent of expert reaches were also recorded for each day. The number of expert reaches and mean correlation coefficients were then compared between VNS, Arch+VNS and control groups in early and late phases.

#### Feature consistency

Several reach features were extracted from each reach attempt: start location (X, Y, Z), end location (X, Y, Z), mean absolute velocity, max absolute velocity, pathlength (length of full trajectory), and reach consistency (defined above in *reach consistency and expert reach*). The distribution of each feature was calculated for the early and late learning phases based on the stimulation group. The distribution of each early-late pair was normalized using the interquartile range of the early phase distribution. The normalized late interquartile range was subtracted from the normalized early range and the difference was defined as ‘delta feature consistency’. A positive delta means that the distribution was more constrained during the late phase compared to the early phase.

### Electrophysiology recording and analysis

#### VNS electrophysiological recording in BF

While mice were either under maintained anesthesia (1.5% isoflurane), or awake in a home cage, VNS was repeatedly delivered while recording from the left basal forebrain (see *optrode implantation*). No behavioral task was performed during recording. Mice received several (10-20) trains of VNS (0.5 s, 30 Hz, 100 µs pulse-width, 0.6 mA), delivered approximately 90 s apart. Data was recorded with Cheetah acquisition software at 30 kHz using a Digital Lynx SX (Neuralynx). TTL pulses were sent from the Master-8 (A.M.P.I.) to the Digital Lynx SX for each pulse of a light or electric stimulation train. In acute experiments, mice were opto-tagged after all VNS trains were delivered to identify cholinergic units.

#### Opto-tagging protocol

While mice were under maintained anesthesia (1.5% isoflurane), recordings were performed in the left BF (see *optrode implantation*). Light was delivered using a class IIIb diode pumped solid state laser (Cobalt) attached to the optrode through a ceramic ferule. Opto-tagging stimulus consisted of several (10-20) trains of 5 mW, 488 nm light delivered just above the BF through a 105 µm diameter fiber optic spaced ~30 s apart. Trains consisted of 10 pulses of light at 20 Hz with a 10 ms pulse duration.

#### Neural classification of BF response

After recording, units were clustered manually using clustering software SpikeSort 3D (Neuralynx) and imported into MATLAB. Isolation distance and L-ratio were used to quantify cluster quality and noise contamination^103^. The start of each stimulation train was identified *post-hoc* using custom scripts (Mathworks) and defined as a trial. The trial window, referred to as the ‘VNS stimulation window,’ was defined as: 1 s baseline before stimulation (−1 to 0 s), VNS delivery (0 to 0.5 s), and 1 s after the end of VNS (0.5 to 1.5 s). Firing rate during the trial window was calculated using a 100 ms moving average, shifted by 1 ms from the start of the trial window to 100 ms after the end of the trial window. Baseline firing rate was defined as the mean firing rate during the baseline period (−1 to 0 s). Units were screened, and any unit with a mean firing rate below 0.5 Hz in anesthetized recordings or below 1 Hz in awake recordings were removed from the pool. ±1 ms around each stimulation pulse was removed to account for electrical noise. Firing rate was converted into a Z-score normalized to the mean firing rate and standard deviation of baseline activity. If a unit’s normalized firing rate was ±2.56 s.d. from the baseline firing rate for >100 ms during VNS delivery (0 to 0.5 s), the unit was defined as VNS responsive. If the change in firing rate was 2.56 s.d. above baseline it was further subclassified as Activated, and if it was 2.56 s.d. below baseline, it was subclassified as Suppressed. A unit that met both criteria was classified based on which occurred first. Peak response, peak delay and response duration were compared between cholinergic and non-cholinergic units in anesthetized mice, and units recorded in awake mice. Peak response refers to the maximum normalized firing rate of VNS activated units after stimulation onset (0 to 1.5 s). Peak response delay refers to the amount of time, in ms, from train onset (0 s) to peak response. Duration refers to the total amount of time that a VNS activated unit had normalized activity <2.56 s.d. above baseline after stimulation onset (0 to 1.5s).

#### Neural classification of opto-tagging in BF

After recording, units were clustered manually using clustering software SpikeSort 3D (Neuralynx). Each pulse was identified *post-hoc* using custom MATLAB scripts (Mathworks) and defined as a trial. Tagged units were identified using a stimulus-associated latency test (SALT^18^). This test compares the distribution of onset time of the first spike recorded during light delivery trials (10 ms) to the onset time of spikes during control windows of similar length (10 ms). Units whose p-value from the SALT test was <0.05 were defined as cholinergic, all other units were defined as non-cholinergic. Firing rates of cholinergic and non-cholinergic units were calculated using the baseline firing rate of the unit from the VNS session.

### Calcium imaging and analysis

#### Data acquisition

Miniscope components and DAQ board were purchased and assembled by Optogenetics and Neural Engineering (ONE) Core according to UCLA miniscope (http://miniscope.org/) V3 guidelines. The objective GRIN lens used were Edmund optics #64-520 and achromatic lens were #45-407. Images were acquired at 30 Hz with Miniscope Control data acquisition package (affiliated with UCLA miniscope). Each imaging session was 15~20 minutes. The calcium signal images were saved as TIFF stacks through USB3.0 port to an SSD hard drive to reduce frame drops. For some home cage sessions, the mice behavior was recorded simultaneously by a LG webcam controlled by MiniscopeControl. For CLARA reach training sessions, behaviors were recorded by the CLARA system and the timing of miniscope frames and behavior camera frames were coordinated.

#### Imaging analysis

##### Neural signal preprocessing

Image stacks that were acquired through the miniscope were motion corrected using MATLAB-based NoRMCorre packages^104^ (https://github.com/flatironinstitute/NoRMCorre). For most sessions, the rigid motion correction module was sufficient to yield good results; in sessions with non-even shifting of the field of view, the non-rigid motion correction module was used. After motion correction, the field of view was cropped and spatially down sampled 4x to reduce the file size for subsequent neural signal extraction. Neural signals were extracted from the images using the MATLAB-based CNMF-E package^105^ (https://github.com/zhoupc/CNMF_E). The results were manually curated to discard non-neuronal ROIs. The resulting C-raw matrix, which was a scaled dF x ROIs, was used for further analysis. For individual ROIs, the time series dF of the whole session was further organized as a 2D matrix dF per trial x trials.

##### Neural classification

To identify VNS responsive neurons, each trial epoch (± 10s around VNS) was mean z-scored to the 3 seconds prior to VNS. Noise response was estimated by calculating average Z-scored dF for individual neurons using a randomly shuffled VNS onset times in the same session (excluding the 3 s window after each VNS onset in the whole session), and bootstrapping across 1000 repeats. For an individual neuron, if the real maximum Z-scored dF in the 3 s window after VNS is higher than the 95% value of the bootstrapped histogram, the neuron was defined as activated by VNS; if the real minimum Z-scored dF in the 3 s window after VNS is lower than the 95% value of randomized z-scored dF, the neuron was defined as suppressed by VNS.

To measure the onset or peak timing for the VNS response, VNS responsive neurons’ dF were Z-scaled with the whole session baseline mean calculated from time points lacking Ca2+ activity^106^ (defined as time points with fluorescence values less than the 0.50 quantile of all fluorescence values). This general-based Z-score process allows dF level above or below 0 before VNS onset each trial and keeps the average dynamics of VNS response more accurately. For individual VNS-activated neurons, the onset of VNS activation was defined as the first time point when the Z-scored dF value reached above 2 s.d. of the mean baseline (the 3 seconds before VNS onset) in the 5 s after VNS onset. For individual VNS suppressed neurons, the onset of VNS suppression was the first time point when the Z-scored dF value reached below 2 s.d. of the mean baseline (the 3 seconds before VNS onset) in the 5 s after VNS onset.

To identify reach-task responsive neurons, a similar trial-based Z-score processes where employed. Trial dFs were aligned by reach max. The baseline was taken as −6 to −3 s before the time of reach max. The response of individual neurons to the task was measured as the maximum and minimum values of the average Z-scored dF in the −800 to −300 ms (reach preparation), −300 to 200 ms (reach), 200 to 1700 ms (outcome), in reference to reach max as time 0. The cut off value for maximum or minimum dF for these time windows were estimated through a similar randomization procedure as above, in which that the same number of reach trials were randomized across the whole session 1000 times. Success trials and failure trials were analyzed separately unless noted otherwise. Several sub-groups of neurons were categorized as: preparation-activated, preparation-suppressed, reach-activated, reach-suppressed, success-activated, success-suppressed, failure-activated and failure-suppressed. Preparation and reach modulated neurons were grouped together as reach modulated neurons in **Fig. 7**. After neural classification, Z-score procedures were used to obtain the average success modulated neurons response to VNS and no-stimulation sessions.

#### Cross-registration of multiple sessions and cross session VNS modulated neurons definition

Due to computational power limits, we chose to process neural activity data from each session and register neurons across sessions. The MATLAB-based CellReg package^107^ (https://github.com/zivlab/CellReg) was used to identify the same neurons from multiple sessions based on spatial footprints of cellular activity (ROIs). Pairs of neurons with correlation coefficient > 0.65 were regarded as the same neurons.

For individual neurons, if a neuron was categorized as success-activated in one of the two training sessions, the neuron was regarded as a success-activated neuron. To look for significant modulation after VNS onset in VNS sessions of these success-activated neurons, the success outcome response of each neuron was aligned to VNS onset and measured in no-stimulation session and VNS session; the difference in response was obtained by subtracting the no-stimulation response from the VNS session response. A difference in response during the 0~3s after VNS onset that was higher or lower than 2.5 s.d. of the mean of baseline (−3 to 0 s before VNS onset) for at least 0.15 s was regarded as significant enhancement or attenuation in the VNS session. To look for significant modulation before VNS onset, the neural response was aligned to reach end. The baseline window was set from −6 to −3 s before reach end. This reach end alignment allows us to evaluate differences in neural representation of reach preparation, execution, and outcome. Differences between the reach representation were compared between the Success VNS and no-stimulation sessions. A difference in response during the −3 to 3 s around reach end that was higher or lower than 2.5 s.d. of the mean of baseline (−6 to −3 s in reference to reach end) for at least 0.15 s was regarded as significant enhancement or attenuation. The onset time of the modulation was defined as the first time point of this enhancement or attenuation.

The control, AChR antagonists, and recovery sessions were processed similarly as VNS in home cage sessions, as described above. In addition, these sessions were temporally down sampled to a 15 Hz frame rate to reduce the file size so that each mouse’s three sessions imaging data could be motion corrected and concatenated together to save the *post-hoc* cross registration step. For VNS-modulated neurons, the VNS activation window was empirically measured and defined as rise onset to peak timing in **Fig. 6g,h** (0.8 to 2.8 s after VNS onset; suppression window as 0.2 to 1.6 s after VNS onset).

### Quantitation and Statistical Analysis

Statistical analyses were conducted using Graphpad, JMP (SAS), or MATLAB (MathWorks). No normality tests were performed but individual data points are plotted to visualize distribution. We used parametric statistics including paired and unpaired Student’s T-tests, and one-way ANOVA with Tukey’s HSD *post-hoc* tests. Two-tailed tests and an alpha cutoff of <0.05were employed unless otherwise stated. We employed a mixed model (Restricted Maximum Likelihood model (REML)) for all learning experiments. REML enables us to test how fixed effects (dependent variables) and known random effects (individual mouse, sex, age) correlate to an outcome variable.

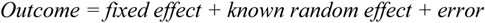

Models were constructed with one or two fixed effects. In cases where there were two fixed effects, we ran full factorial models. To determine early and late phases of learning, a Weibull growth curve was applied to Sham VNS learning data. The formula was:

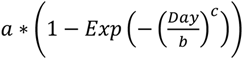

a = asymptote, b = inflection point, c = growth rate. The AICc = 1295.90 and the R^2^ = 0.21. We used the asymptote of the control cohort (55.50% ± 6.8%) to represent the plateau in success rate. The late learning phase was selected based on the first training day that exceeded the lower 95^th^ percentile of the asymptote (42.15%), which was day 4. We thus called late learning days 5-14 and early learning days 1-4.

## Notes

### Competing Interest Statement

The authors have declared no competing interest.

### Summary of Updates

Final version submitted to the journal, updates to grammar and figures.

